# RGG motif-containing Scd6/LSM14A proteins regulate the translation of specific mRNAs in response to hydroxyurea-induced genotoxic stress

**DOI:** 10.1101/2022.07.12.499753

**Authors:** Gayatri Mohanan, Raju Roy, Hélène Malka-Mahieu, Swati Lamba, Lucilla Fabbri, Anusmita Biswas, Sylvain Martineau, Céline M. Labbé, Stéphan Vagner, Purusharth I Rajyaguru

## Abstract

Genotoxic stress response (GSR) mediated by mRNA translation and decay regulation remains poorly explored. Here, we identify a unique role of yeast RGG-motif protein Scd6 and its human ortholog LSM14A in mRNA translation control upon hydroxyurea (HU)-mediated GSR. Scd6/LSM14A, but not all tested RGG-containing proteins, localize to HU-induced cytoplasmic puncta in an RGG-dependent manner. The absence of Scd6 increases HU tolerance but sensitizes the cells to HU upon overexpression of SRS2, a known dampener of the DNA-damage response. Scd6 binds *SRS2* mRNA to repress its translation in cytoplasmic granules upon HU stress. Scd6-SRS2 interaction is modulated by arginine methylation (AM) and the LSm-domain, which acts as a *cis*-regulator of Scd6 AM. Polysome-profiling experiments indicate that LSM14A regulates the translation of NHEJ factor mRNAs such as *LIG4* (DNL4 homolog) and *RTEL1* (SRS2 functional homolog), and the NHEJ activity in response to HU. Overall, this report unveils the role of AM and Scd6/LSM14A in the GSR by determining the translation status of specific mRNAs.

## Introduction

DNA damage caused by cellular conditions and/or exogenous factors leads to genomic instability. Replication errors, defective DNA damage repair, and nucleotide misincorporation represent cellular events leading to DNA damage ^1^. Exogenous factors causing DNA damage include exposure to genotoxins such as hydroxyurea, cisplatin, etc. Living systems employ strategies to regulate gene expression and adapt the cellular proteome to mount an effective genotoxic stress response (GSR). Even if the contribution of gene expression to GSR is overall well documented, changes in the fate of cytoplasmic mRNAs in response to GSR remain poorly explored^2^.

RNA binding proteins (RBPs) play a crucial role in determining the functional states of mRNAs. Several classes of RNA binding domains, such as RNA Recognition Motif (RRM), KH-domain, Zn-finger motif and PUMILIO, have been reported to play a role in post-transcriptional gene expression by changing the fate of specific subsets of mRNAs. RGG motif-containing proteins are the second largest class of RBPs^3,4^. RGG-motifs are characterized by RGG-/RGX repeats that impart properties of low-complexity sequences^3^. Consistently, these sequences contribute towards the assembly of ribonucleoprotein (RNP) condensates (*i.e.,* RNA granules) by undergoing liquid-liquid phase separation^5^. In yeast, processing bodies (P-bodies or PB) and stress granules (SG) are the major cytoplasmic mRNPs formed in response to several physiological cues. They contain several RNAs and RBPs. PBs are reported to be sites of mRNA decay, whereas SGs are implicated in mRNA storage and repression^6^. *S. cerevisiae* Sbp1 and Scd6 are two RBPs with RGG-motif sequences that target eIF4G to repress translation^7^. Scd6 is a known P-body and stress granule resident protein in response to glucose deprivation and oxidative stress^8^. The RGG motif of these proteins is important for localization to RNA granules such as P bodies, interaction with eIF4G, and consequent translation repression activity^8,9^. Interestingly, the RGG-motif of Sbp1 has recently been implicated in P-body disassembly, suggesting that RGG-motifs can promote both assembly and disassembly of RNA granules^10^. LSM14A, the human ortholog of Scd6, is a granule-resident protein that has also been implicated in translational control^11^. LSM14A contains two RGG motifs as compared to a single RGG motif in Scd6^12^. LSM14A plays a crucial role in forming the mRNA silencing complex via its association with DDX6^13^. Although xRAP55 (Xenopus ortholog of LSM14/Scd6) has been reported to repress translation^14^, the direct role of human LSM14 in translational repression remains to be demonstrated. Interestingly, LSM14A has been reported as a sensor of viral nucleic acid, which plays a crucial role in antiviral response^15^.

The role of LCS-containing proteins in GSR remains poorly explored. Even though the localization of Scd6 to puncta upon HU stress is known^16^, the functional relevance of the Scd6 family of proteins with RGG motif in genotoxic stress response remains unexplored. In this report, we identify the contribution of Scd6/LSM14A in regulating translation of specific mRNAs following HU-mediated genotoxic stress. We further provide mechanistic insight into arginine methylation-mediated regulation of Scd6-SRS2 interaction during HU stress.

## Results

### Scd6 and Sbp1 localize to granules in response to HU stress

To assess the role of RGG-motif proteins in hydroxyurea-mediated stress response, we tested the localization of five RGG-motif-containing RNA-binding proteins (Scd6, Sbp1, Npl3, Psp2 and Gbp2) upon HU stress (Figure S1A). Scd6, Sbp1 and Npl3 are known to be involved in translation repression activity mediated by binding to eIF4G1^7^. Psp2 has been implicated in promoting the translation of specific mRNA in autophagy response^17^. Gbp2 is mainly involved in mRNA export and processing; however, recently, it has been implicated in translation repression^18^. Npl3 and Gbp2 are shuttling proteins, primarily localised in the nucleus and have known connections to cellular stress response^19,20^. Initial time course analysis revealed that Scd6 localized to distinct cytoplasmic puncta upon 45 min treatment of 0.2M HU (Figure 1A & 1C), whereas Sbp1 localised to foci upon 60 min of 0.2M HU treatment (Figure 1B & 1D). No significant change in localization was observed for GFP-tagged Npl3, Gbp2, and Psp2, even after 120 minutes of HU treatment (Figure S1B), indicating that localization to puncta upon HU stress is not a general response shared by RGG-motif proteins but a specific feature of Scd6 and Sbp1. Localization of both Scd6 and Sbp1 to granules is not due to increased protein levels upon HU exposure, as indicated by western blot analysis (Figure S1C, S1D). Of note, due to technical difficulty associated with detecting Scd6-GFP using anti-GFP antibody, levels of endogenously tagged Scd6-myc were quantified upon HU treatment. Colocalization studies indicated that 91% of Sbp1-GFP foci colocalized with Scd6-mCherry foci upon 60 minutes of HU exposure (Figures S1A and S1B). Localization of Scd6 and Sbp1 to puncta was visible after 45 and 60 min, respectively, indicating that Scd6 localization was relatively sooner than Sbp1. We, therefore, wondered if Scd6 promoted Sbp1 localization to puncta and observed that Sbp1 localization to puncta upon HU-stress was defective in *Δscd6* with no significant decrease in protein levels as indicated by GFP intensity measurements (Figures S1C and S1D). However, Scd6 localization to puncta was unaffected by *Δsbp1* (Figures S1E and S1F). Our results provide evidence for specific and coordinated localization of Scd6 and Sbp1 to HU- induced puncta.

**Figure 1.**
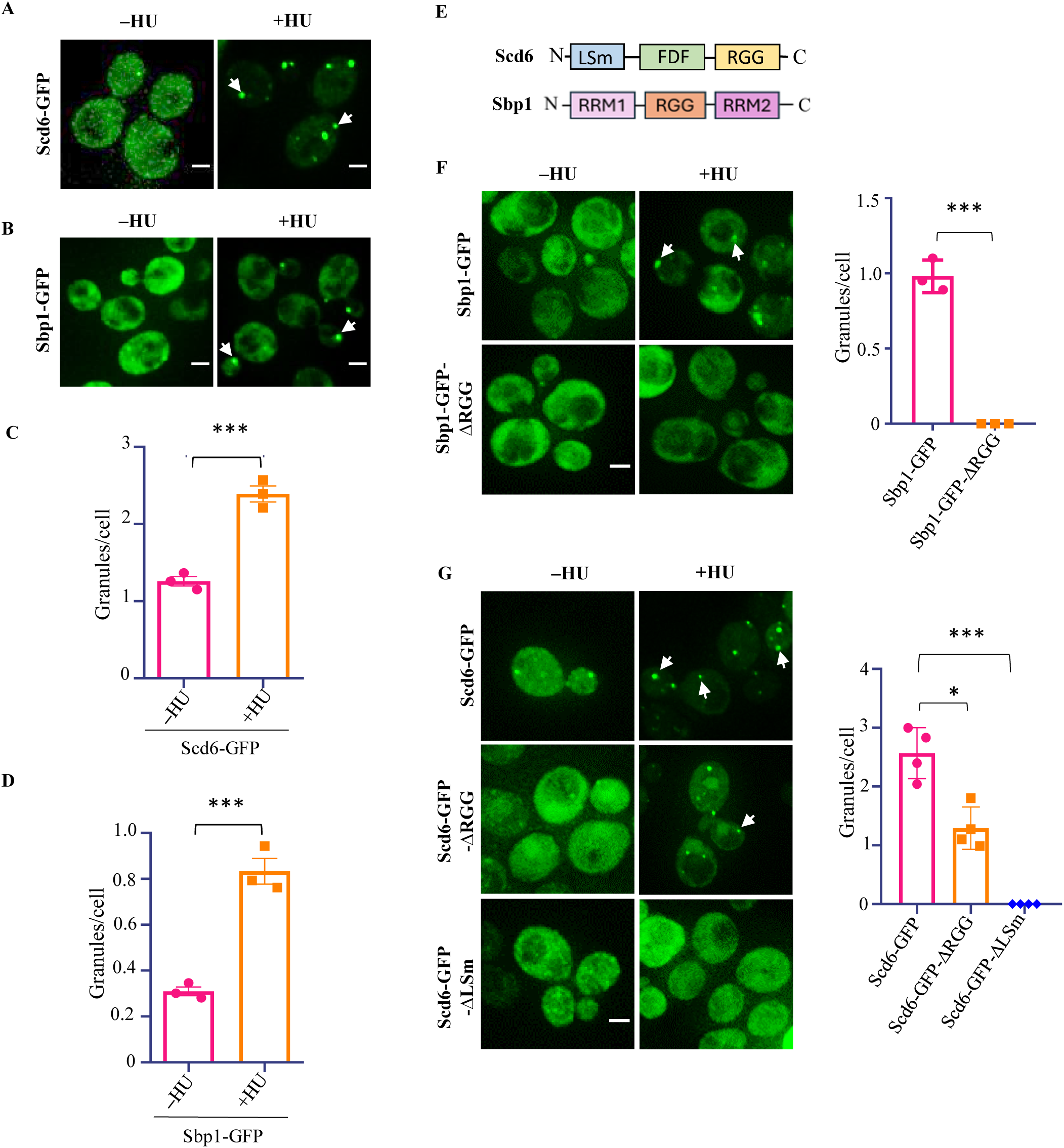
Both LSm domain and RGG-motif are important for the localization of Scd6 to granules in response to HU. (A) and (B) Live cell imaging showing localization of plasmid-borne (*CEN*) Scd6-GFP and endogenously tagged Sbp1-GFP to granules upon 200 mM HU treatment (45 minutes treatment for Scd6-GFP and 60 minutes for Sbp1-GFP). (C) and (D) Quantification of granules as granules per cell (E) Schematic representation showing domain organization of Scd6 and Sbp1. (F) and (G) Live cell imaging showing localization of Sbp1 and Scd6 and their domain deletion mutants upon HU treatment and their quantification. n=3; ∼300 cells were counted for analysis for all experiments. White arrows indicate cytoplasmic foci. Statistical significance was calculated using unpaired t-test. Error bars indicate the standard error of mean. Scale bar denotes 2 μm.

### Both the LSm domain and RGG-motif are important for localization to puncta

Scd6 and Sbp1 are multidomain proteins containing RGG motifs at the C-terminal and in the middle, respectively (Figure 1E). In addition, Scd6 has an N-terminal LSm domain and a middle FDF domain (Figure 1E, top). This domain organization is conserved across all Scd6 orthologs^12^. The RGG domain of Sbp1 is flanked by two RRM domains, RRM1 and RRM2 (Figure 1E, bottom). RGG motifs constitute low-complexity sequences implicated in the assembly of RNA granules. Since the translation repression activity of both Scd6 and Sbp1 was mediated through their RGG domain^8,9^, we tested the localization of the RGG domain deletion mutant of both Scd6 and Sbp1. We observed that the localization of the Scd6ΔRGG mutant to puncta decreased by nearly 50% (Figures 1G) whereas the localization of the Scd6ΔLSm mutant to puncta was almost completely abrogated (Figure 1G, bottom). Interestingly, deletion of the Sbp1 RGG motif (Sbp1ΔRGG) caused complete cessation of Sbp1 localization to puncta (Figure 1F). These observations suggest that both the LSm domain (for Scd6) and RGG-motif (for both Scd6 and Sbp1), which have been implicated in the translation repression activity of these proteins, are important for their localization to the puncta upon HU stress. Overall, our results argue for the role of Scd6 and Sbp1 in HU-mediated stress response.

### HU-induced puncta are dynamic and sensitive to cycloheximide

To investigate whether mRNAs were present in the Scd6 and Sbp1 puncta formed upon HU treatment, we treated cells with cycloheximide (CHX), which locks the mRNA on polysome, thereby reducing its availability for granule assembly and maintenance. When HU-treated cells were subjected to 0.1 mg/ml of CHX for 5 minutes at 30 °C (Figure S2E), there was a substantial decrease in both Scd6-GFP and Sbp1-GFP granules (Figures S2F). This suggests that mRNAs are present in HU-induced Scd6 and Sbp1 puncta.

RNA granules disassemble rapidly upon removal of stress^21^. The reversible nature of RNA granules is essential as it allows the return of mRNAs and various RBPs to the cytoplasm. To examine if the HU-dependent formation of Scd6 and Sbp1 granules is reversible, we performed the experiments described earlier but followed by a recovery period (Figure S2G). The HU-treated cells were resuspended and grown in HU-free media during the recovery period. After 75 minutes of recovery for Scd6 and 90 minutes of recovery for Sbp1, both Scd6 and Sbp1 puncta decreased significantly (Figure S2H). Based on these observations, we conclude that the formation of Scd6 and Sbp1 granules in response to HU is reversible. Since CHX experiments indicated that these granules contain mRNAs, the localization of Scd6 and Sbp1 to these puncta could be involved in regulating the fate of specific mRNAs in response to HU.

### Deletion of SCD6 increases sensitivity to HU upon SRS2 overexpression

To test the role of Scd6 and Sbp1 in the response to HU, we measured the growth kinetics of *Δscd6* and *Δsbp1* strains upon HU treatment. We observed that both strains grew marginally but significantly better than the wild-type (WT) strain in the presence of HU reflected by lesser lag-time required to achieve the maximum growth rate (Figures 2A and 2B). Consistent with this observation, we observed an increase in the CFUs of both *Δsbp1* and *Δscd6* strains on HU-containing media as compared to the WT strain (Figure 2C). These observations indicate that Scd6 and Sbp1 contribute to HU-stress response pathway(s).

**Figure 2.**
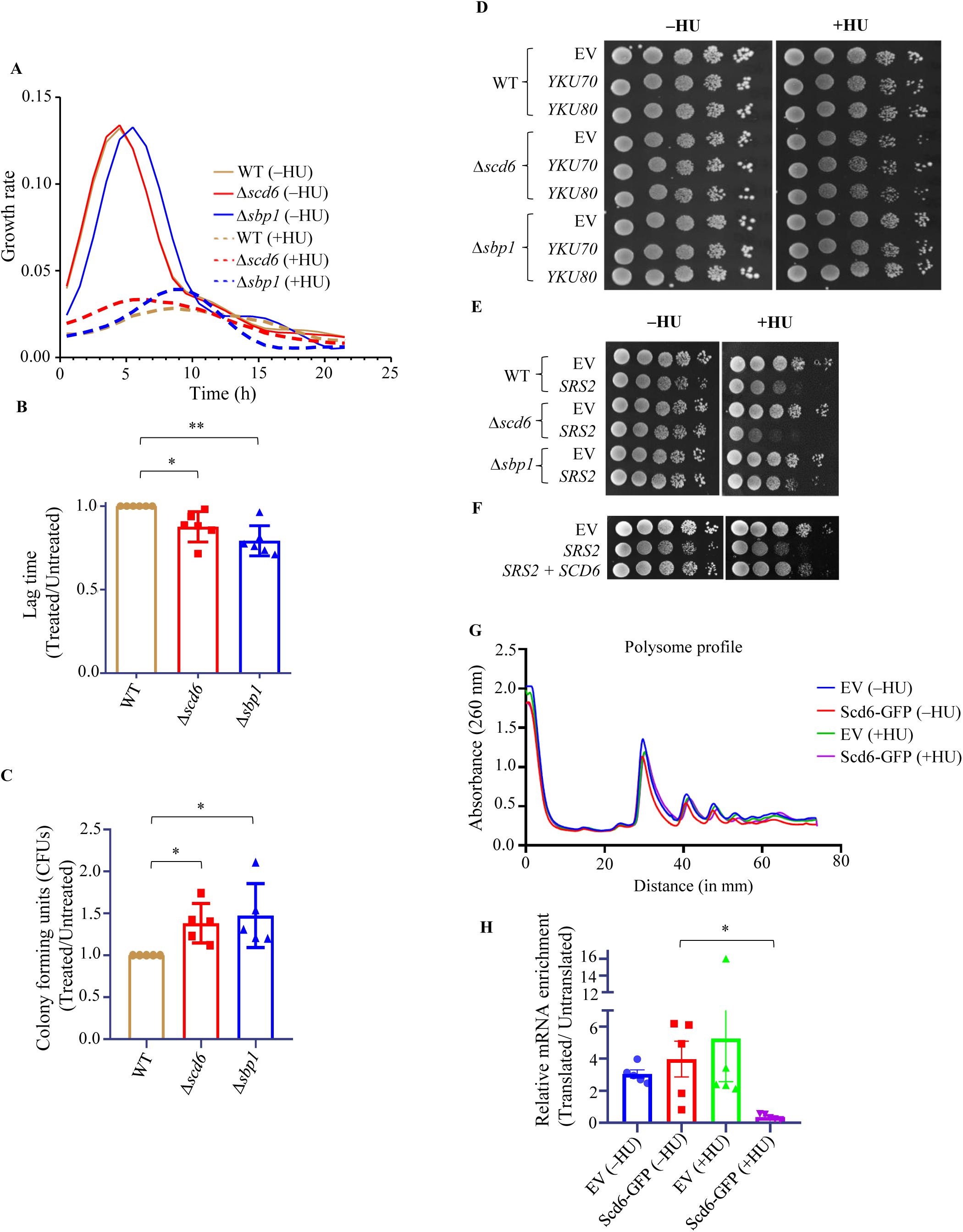
Scd6 modulates cellular tolerance to HU by translation repression. (A) Representative growth rate curve comparing WT and deletion strains in untreated, and 200 mM HU treated media. (B) Quantification of growth rate in terms of lag time, plotted as change w.r.t. WT (n=6). (C) Quantification of plating assay, plotted as CFUs normalized against WT (n=5). 100 ul of 10^-4^ dilution cell suspension were plated on control (−HU) or 100 mM HU-containing YEPD plates. (D) Spot assay of WT, *Δscd6,* and *Δsbp1* transformed with either CEN empty vector (EV*), YKU70 or YKU80* plasmids and spotted on Control and 100 mM HU containing Leucine dropout media and 2% glucose. (E) Spot assay of WT and deletion strains either carrying EV, or SRS2 on CEN plasmid, spotted on Uracil dropout agar plate and 2% glucose, in the presence or absence of 100 mM HU. The plates were incubated in 30°C. (F) Spot assay with WT either carrying EV, or SCD6 on 2µ plasmid, along with SRS2 on CEN plasmid, spotted on Uracil-Leucine dropout agar plate and 2% glucose, in the presence or absence of 100 mM HU. (G) Representative polysome profiles of 200 mM HU treated or Control cells expressing Scd6-GFP on a 2µ plasmid (or EV) (H) Quantification of *SRS2* mRNA in the polysome fractions plotted as relative log2-Fold change ratio of Translated/Untranslated fractions (n=5) using SRS2 specific primer normalized to total mRNA and PGK1 (internal control). Error bars indicate standard error of mean and statistical significance was calculated using unpaired t-test.

HU exposure leads to reduced dNTP availability by inhibiting RNR activity. This can lead to replication fork collapse, resulting in DNA double-stranded break^22^. Since Scd6 and Sbp1 are known translation repressor proteins, we wondered if it could modulate the role of certain factors implicated in genotoxic stress response. We observed a marginal but significant decrease in Rnr3 protein (a direct target of HU) levels upon HU treatment in *Δscd6* and *Δsbp1* strain compared to WT (Figure S3A). The protein levels of Rad53, which is an upstream sensor kinase involved in the GSR signalling pathway, were not significantly changed across the strains upon HU treatment (Figure S3B). Since *Δscd6* and *Δsbp1* showed tolerance to chronic HU exposure, and Rnr3 protein levels were slightly decreasing, we hypothesized that the deletion strains were adapting more efficiently to HU-induced genotoxic stress than the WT strain. We therefore tested if Scd6 or Sbp1 affected the response of cells to HU treatment upon overexpression of several factors known to be involved in the repair of damaged DNA such as YKU70, YKU80 and SRS2. The levels of SRS2, a 3’-5’ DNA helicase involved in regulating homologous recombination (HR)^23^ are tightly regulated, and overexpression of SRS2 causes growth defects and increases sensitivity to genotoxic stress^24^. Excess SRS2 causes the formation of toxic intermediates upon checkpoint activation and initiation of HR-mediated repair ^25^. A recent study also showed that SRS2 dampens the DNA damage checkpoint, leading to genotoxin tolerance ^26^. We observed that the absence of Scd6 or Sbp1 did not affect the cell growth upon overexpression of YKU70 and YKU80 (Figure 2D). However, overexpression of SRS2 compromised cell growth in the presence of HU, this growth defect being stronger in the *Δscd6* background than in the Δ*sbp1* background (Figure 2E, S3C). In addition, overexpression of SCD6 completely rescued the growth defect caused by SRS2 overexpression in the presence of HU (Figure 2F). Altogether, these results provided an important insight into the role of Scd6 in the HU-mediated stress response, indicating that Scd6 regulated the expression of SRS2 in an HU-dependent manner. Given the role of Scd6 as a translation repressor, we hypothesized that Scd6 might regulate the translation of the *SRS2* mRNA.

### SCD6 overexpression reduces *SRS2* mRNA association with translating ribosomes upon HU stress

To test the role of Scd6 in mRNA translation upon HU stress, we overexpressed Scd6-GFP in a 2µ plasmid under its own promoter and looked at its effect on global translation using polysome profiling. Scd6 overexpression did not cause global translation defects, nor did HU treatment (Figure 2G). To test the role of Scd6 on *SRS2* mRNA translation upon HU stress, we isolated RNA from the sucrose gradient fractions followed by qRT-PCR to estimate the amount of *SRS2* mRNA in polysome fractions. We found that, upon Scd6 overexpression in HU-treated cells, there is a significant decrease in the amount of *SRS2* mRNA present in polysome fractions (normalized by SRS2 mRNA present in untranslated fraction) as compared to in untreated cells (Figure 2H). The association with polysomes of the *DNL4* (DNA ligase IV) and *RNR4* (Ribonucleotide Reductase 4), which are known to be involved in HU-induced DNA damage response, did not change under the same conditions (Figure S3D). These results indicate that Scd6 specifically represses the translation of the *SRS2* mRNA upon HU stress.

### Scd6 interacts with the *SRS2* mRNA

To investigate the possible HU-dependent association of Scd6 to the *SRS2* mRNA, we performed RNA immunoprecipitation (RIP) in cell lysates of cells expressing Scd6-GFP in the presence of HU (*i.e.* in the same conditions as the polysome profiling experiment in Figure2G) (Figure 3A and 3B). The amounts of *SRS2* mRNA expression were quantified in the Scd6 pull-down fractions immunoprecipitates (IP) and normalized with total *SRS2* mRNA and *PGK1* control (using PGK1-specific primers). We observed a significant enrichment of *SRS2* mRNA in the Scd6 IP from HU-treated cell lysate (Figure 3C). This enrichment was not visible when the experiment was performed in cells expressing ΔLSm or ΔRGG mutants of Scd6, indicating a role of these protein domains in the Scd6-*SRS2* mRNA interaction. Transcripts encoding two key DNA damage response proteins, DNL4 (the yeast homolog of Ligase IV) and RNR4, were also tested. The *DNL4* mRNA, but not the *RNR4* mRNA, was significantly enriched in the Scd6 IP when cells were exposed to HU (Figure S4A), in a similar manner as the *SRS2* mRNA. According to the polysome profiling results, the *DNL4* mRNA was however not translationally repressed by Scd6 (Figure S3D). We believe that this enrichment could be a result of indirect interaction mediated by another protein in the complex. Alternatively, it is possible that Scd6 could affect the stability of the *DNL4* transcript.

**Figure 3.**
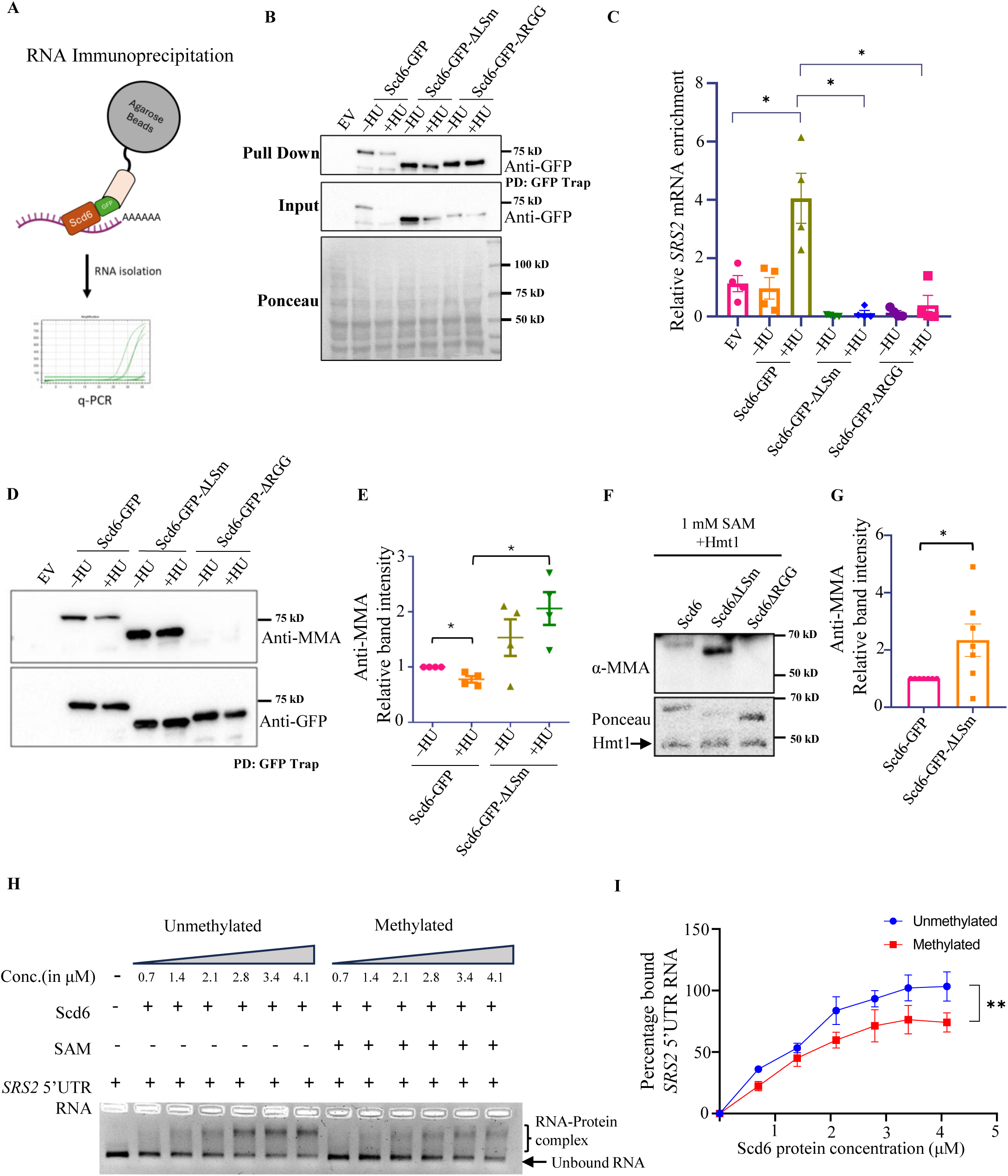
Scd6 binds to *SRS2 mRNA* upon HU treatment. (A) Approach for RNA immunoprecipitation (RIP) using GFP-tagged Scd6 (expressed in a 2 µ plasmid, under its own promoter) in untreated (–HU) and 200 mM HU treatment. (B) Representative western blot showing pull down of Scd6-GFP WT and domain deletion mutants used in RIP (C) Quantification of *SRS2* mRNA enriched in the pull-down fraction, plotted as relative log_2_-Fold w.r.t total mRNA (n=4). (D) Western blot showing arginine-mono-methylation of Scd6-GFP WT and domain deletion mutants upon HU stress which is quantified in (E) n=4. (F) Western blot for *in vitro* arginine-mono-methylation of recombinant WT and domain deletion mutants of Scd6 incubated with equimolar concentration of recombinant Hmt1 and 1 mM S-adenosyl methionine (SAM) for 120 mins at 37 °C. (G) Quantification of relative *in vitro* mono- methylation as seen in F (n=7). Statistical significance was calculated using unpaired t-test (H) Ethidium Bromide (EtBr) stained agarose gel for electromobility shift assay (EMSA) with increasing concentrations of recombinant Scd6 with or without *in vitro* methylation incubated with 200-mer 5’UTR fragment of *SRS2 mRNA* (0.94 μM RNA) (I) Quantification of percentage bound *SRS2 mRNA* fragment as a function of protein concentration (n=3). Statistical significance calculated by paired t-test. Error bars indicate standard error of mean.

### An HU-dependent decrease in Scd6 methylation increases Scd6-*SRS2* mRNA interaction

Scd6 is arginine methylated at RGG-motif, which regulates repression activity^9^. However, the direct role of arginine methylation in Scd6 interaction with RNA is not known. We checked if a change in the methylation status of Scd6 could mediate the increased interaction of Scd6 with the *SRS2* mRNA upon HU stress. A western blot using an antibody against mono-methylated arginine revealed a signal for Scd6 which was absent, as expected, for the RGG deletion mutant. We observed a slight but consistent decrease in Scd6 methylation upon HU stress. Interestingly, this was not the case with the Scd6 mutant lacking the LSm domain, whose methylation levels were significantly more than the WT Scd6 in the presence of HU (Figure 3D and 3E). This suggested that the LSm domain could negatively regulate the methylation status of Scd6, thereby promoting RNA binding upon HU stress which is consistent with the observations that the LSm domain or the RGG domain is required for RNA interaction (Figure 3C). To test if the absence of the Lsm mutant could directly affect methylation status, purified Scd6 and Lsm domain-deletion mutants were incubated with purified Hmt1 (the methyltransferase known to methylate Scd6) (Figure S4B). We observed increased methylation of the Lsm domain deletion mutant as compared to the wild type (Figure 3F-G), suggesting A direct role of the Lsm domain in the cis-regulation of methylation. Since HU treatment reduced Scd6 methylation (Figure 3D) and increased Scd6 interaction with the *SRS2* mRNA (Figure 3C), we hypothesized that arginine methylation could inhibit Scd6-*SRS2* mRNA interaction *in vitro*. We found that methylated Scd6 interacted less efficiently than unmethylated Scd6 with an *in vitro* transcribed RNA containing the 5’UTR of the *SRS2* mRNA (Figure 3H and 3I, S4C) showing that the HU-dependent regulation of Scd6 methylation is important for RNA binding.

### The *SRS2* mRNA associates with Scd6-containing granules in HU-treated cells

Our results suggest that the HU stress increases the localization of Scd6 to granules and its interaction with the *SRS2* mRNA. To check if this interaction occurred in granules, we used two approaches. The first one involved the quantification of *SRS2* mRNA (RT-qPCR) and Scd6 protein (WB) in the cytoplasmic (lighter) and granule-enriched (heavier) fractions from lysates of cells treated with HU (Figure S5A)^27^. As previously reported for yeast stress granule cores, Scd6-containing granules were resistant to NaCl and EDTA treatment, indicating that these granules are stabilized by hydrophobic interactions. RNase treatment did not perturb these structures either, indicating that protein-protein interactions play a major role in the integrity of the granules once the core is formed. As expected, 1% SDS treatment caused a complete loss of Scd6 signal from the pellet fraction and appeared in the soluble fraction, confirming that these are not Scd6 protein aggregates but higher order mRNA-protein complexes harbouring Scd6 (Figure S5B). We found a significant enrichment of the *SRS2* mRNA in granules-enriched fractions from HU-treated cells expressing Scd6-GFP. The enrichment of the *SRS2* mRNA in granules-enriched fractions was, however, significantly less in HU-treated cells expressing the LSm domain-deletion mutant of Scd6 (Figure 4A), which is itself less enriched in granule-enriched fractions than the WT Scd6-GFP protein.

**Figure 4.**
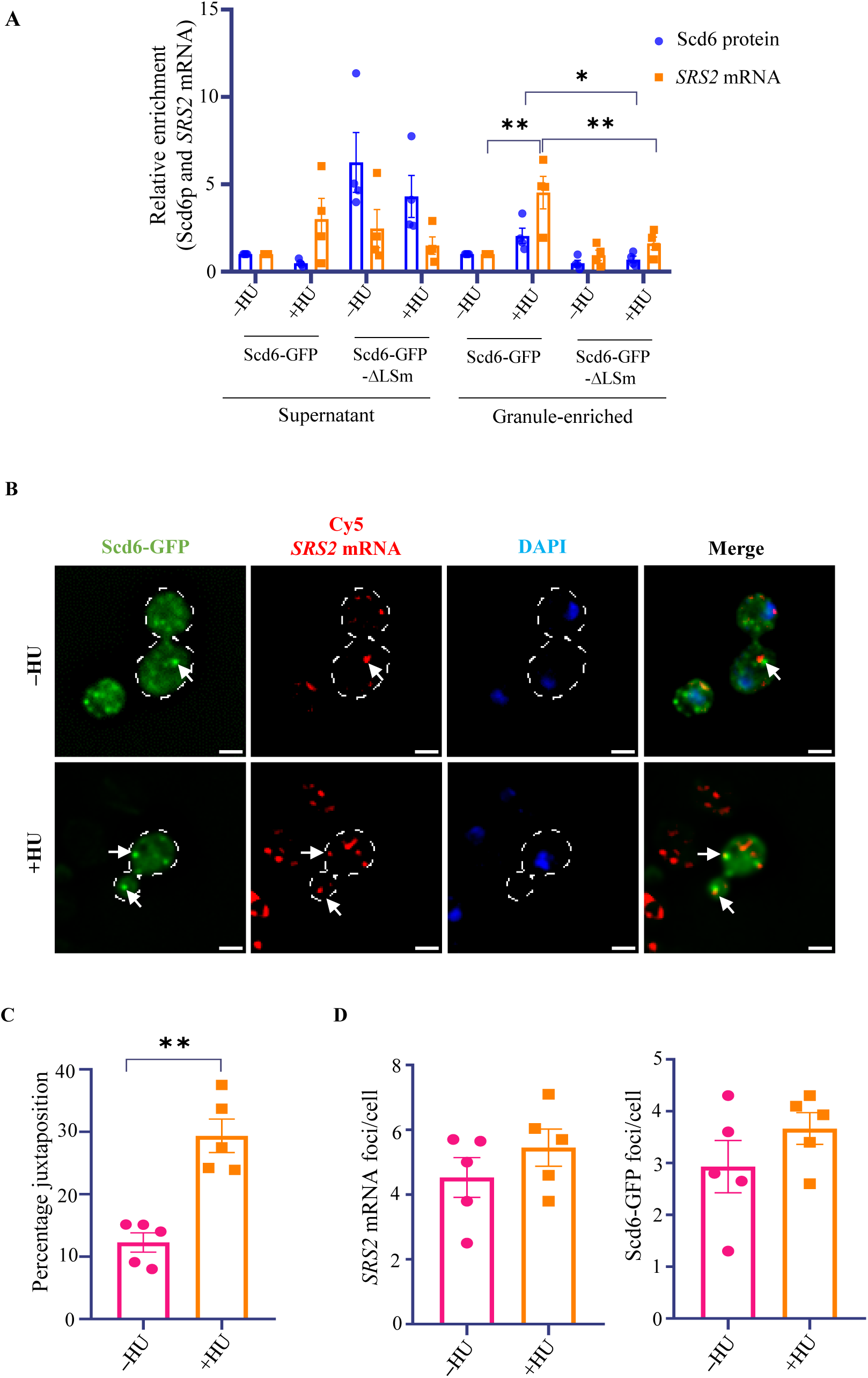
*SRS2* mRNA localizes to granule enriched fraction upon HU stress. (A) Relative enrichment of Scd6 protein and *SRS2* mRNA in heavier granule enriched fraction (n=4). Statistical significance was calculated using One-Way Annova. (B) smFISH showing localization of *SRS2* mRNA and Scd6GFP in control and HU-treated cells. White arrows indicate juxtapositioned *SRS2* and Scd6 foci. Scale bar denotes 2 μm. (C) Quantification of the percentage of *SRS2* mRNA juxtapositioned with Scd6 granules normalized to total Scd6 granules (n=5, ∼100 cells counted). (D) Quantification of number of *SRS2* and Scd6 foci formed upon HU stress compared to unstressed control for the same experiment. Statistical significance was calculated by paired t- test. Error bars indicate standard error of mean.

The second approach used smFISH (single-molecule fluorescence *in situ* hybridization) to detect the localization of the *SRS2* mRNA using Cy5-labelled secondary probes with affinity to SRS2 specific oligo pool. Upon HU treatment, there was a significantly increased juxtaposition of *SRS2* mRNA foci with Scd6 granules (Figures 4B and 4C), even though the total number of *SRS2* mRNA or Scd6-GFP protein foci did not change significantly (Figure 4D). These results indicate that the Scd6 protein and the *SRS2* mRNA are localize to granules upon HU stress.

### LSM14A localizes to puncta upon HU treatment in an RGG-motif-dependent manner and modulates translation

To analyze whether the observed role of Scd6 in genotoxic stress response was conserved in humans, we focused on LSM14A, the human ortholog of Scd6, which is known to localize to P-bodies and stress granules^11^. A direct role of LSM14A in repressing translation remains to be demonstrated. Like Scd6, LSM14A is a modular protein (Figure 5A) with two RGG domains. Using live cell imaging, we found that LSM14A localized to puncta upon HU treatment in an RGG-motif-dependent manner, (Figures 5B and 5C), indicating that it is a conserved feature of the Scd6 family of proteins in yeast and human. To examine the role of LSM14A in regulating mRNA translation in response to HU treatment, we performed polysome profiling experiments (Figure 5D). RNAs isolated from translating (heavy) and non-translating (light) fractions of wild type and LSM14A knockdown (siRNA) cells were sequenced to identify mRNAs whose association with polysomes was perturbed by LSM14A (Supplementary Table S1). Several mRNAs were identified to be differentially regulated. To address the role of LSM14 in the HU response, a similar analysis was carried out for LSM14A-depleted cells treated with HU. Again, many mRNAs were identified to be differentially regulated at the translation level, highlighting an important role of LSM14A in translation control under normal conditions and genotoxic stress (Figure 5E).

**Figure 5.**
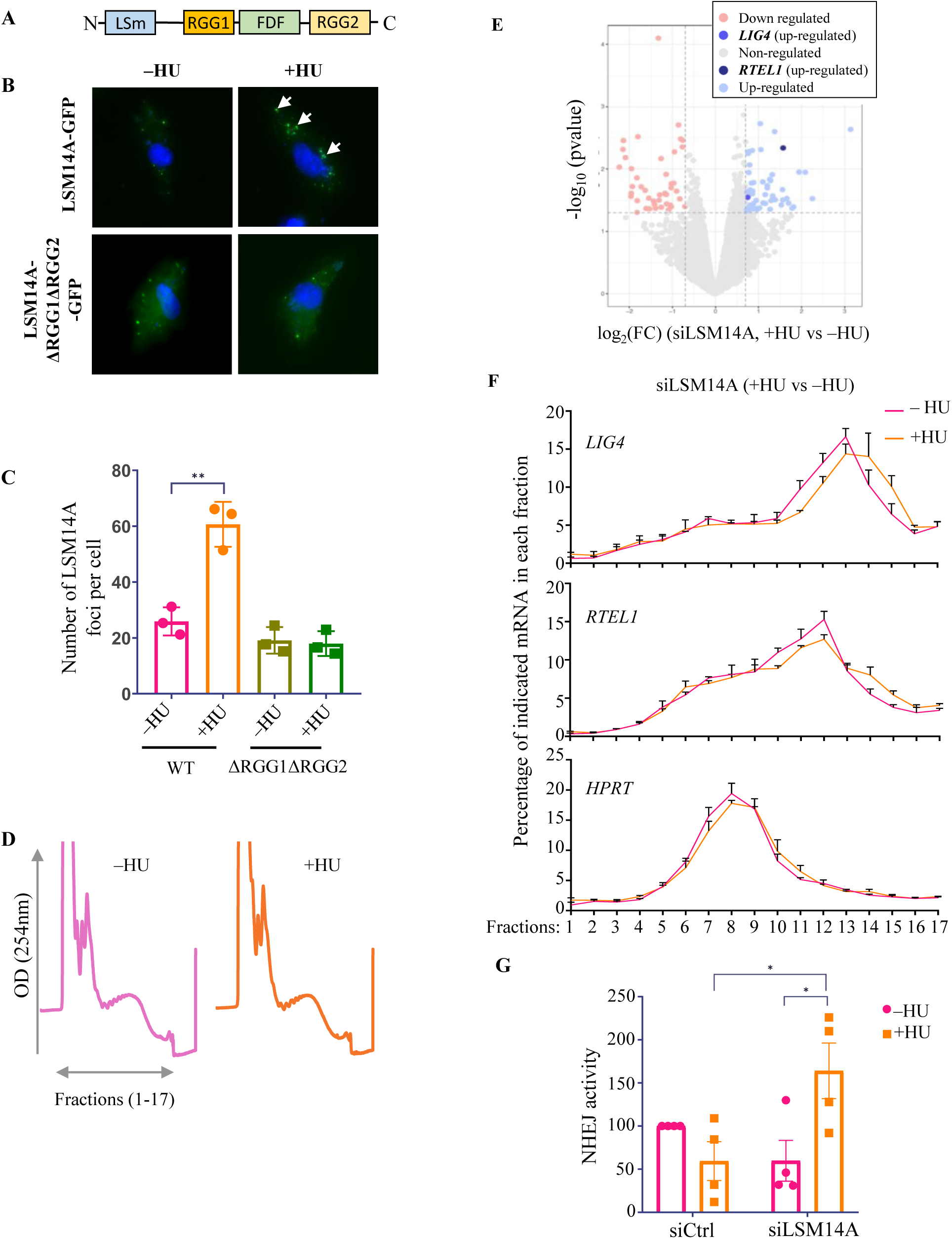
LSM14a localizes to puncta upon HU in an RGG-motif dependent manner and regulates mRNA translation. (A) Domain organization of LSM14A. LSM14A contains a N-terminal LSm domain, a FDF motif and two RGG motifs flanking the FDF motif. (B) Localization of LSM14A WT and RGG-deletion mutants upon HU treatment in RPE-1 cells. (C) Quantitation of localization of LSM14A WT and RGG-deletion mutants as depicted in B. Statistical significance was calculated using paired t-test. Error bars indicate standard error of mean. (D). Polysome profiles of siLSM14A-transfected A2058 cells treated or not with HU. OD: Optical density. (E). Volcano plots report the log_2_ fold change −log_2_(FC)- on the x-axis and the minus log_10_ of the p-value –log_10_(p-value) on the y-axis obtained from the comparison of translation efficiencies in the +HU vs –HU treated A2058 cells (Table S1). Blue dots highlight the translationally up- regulated mRNAs while red dots highlight the translationally down-regulated mRNAs . (F) Validation of *LIG4* and *RTEL1* mRNA in each polysome fraction comparing siLSM14A untreated (–HU) condition with siLSM14A HU treated condition. Distribution of control *HPRT* mRNA in the polysome fractions is also shown. N=3 replicates. (G) Quantification of plasmid integration efficiencies in A375 cells transfected with the indicated siRNAs and treated with or without HU. Data were normalized to control transfection without HU treatment, which was set to 100%. Statistical significance of the experimental data was determined using Two-Way ANOVA (mean ± SEM, n=4).

We observed that the association of transcripts encoding DNA damage repair proteins, including Ligase IV (NHEJ pathway protein, DNL4 yeast homolog) and RTEL1 (the functional homolog of SRS2), with translating fractions increased upon HU stress in cells depleted for LSM14A (Figure 5E). RT-qPCR analysis of RNAs extracted from each fraction of the sucrose gradient confirmed this regulation since we observed a slight shift in the distribution of the *Ligase IV* and *RTEL1* mRNAs, but not the *HPRT* mRNA used as a negative control, towards heavy polysome fractions upon HU treatment (Figure 5F). To examine if the translation regulation of genes involved in NHEJ is associated with a change in NHEJ activity, we monitored NHEJ activity using a random plasmid integration assay. In this assay, linearized DNA with a neomycin selection gene is transfected and the efficiency of random chromosomal integration of the plasmid DNA by NHEJ is measured by colony formation in G418-containing media. We observed that LSM14A knockdown significantly increases the NHEJ activity upon HU stress (Figure 5G), indicating that LSM14A-mediated translation regulation is associated with the DNA damage response to HU stress.

## Discussion

In this report, we have explored and identified a conserved role of RGG motif-containing Scd6 family proteins in genotoxic stress response to HU. Several observations support this conclusion: (i) Sbp1, Scd6 and its human homolog LSM14A localize to puncta upon HU-induced genotoxic stress in an RGG-motif dependent manner (Figure 1 and 5), (ii) Scd6 represses *SRS2* mRNA translation upon HU stress (Figure 2), (iii) Scd6 binds to *SRS2* mRNA *in vivo* upon HU stress (Figure 3), (iv) Arginine methylation of Scd6 is reduced upon HU stress (Figure 3), (v) LSm domain deletion mutant is hypermethylated and defective in binding *SRS2* mRNA (Figure 3), (vi) Methylation reduces Scd6 protein interaction with *SRS2* mRNA (Figure 3), (vii) HU-induced granules are sites of *SRS2* mRNA enrichment and repression (Figure 4) (viii) LSM14A regulates translation of multiple transcripts upon HU stress including *LIG4* and *RTEL1* (Figure 5), and (ix) LSM14A knockdown increases NHEJ activity upon HU stress (Figure 5). These results identify an exciting new role of Scd6 family proteins in the genotoxic stress response.

HU is a well-known genotoxin that causes DNA damage by depleting dNTP pools, leading to replication fork arrest eventually causing DNA double-strand breaks ^28–30^. Genome-wide studies have reported changes in the localization of numerous proteins in response to genotoxic stress^16^; however, barring a handful of studies, the functional relevance and mechanistic basis of altered localization have not been explored. RNA granules are sites of regulation of mRNA translation and decay. Localization of cytoplasmic RNA binding proteins Sbp1, Scd6, and LSM14A to puncta upon HU treatment suggests that such localization change could lead to regulation of mRNA fate upon HU stress.

Nuclear shuttling proteins such as Gbp2 and Npl3 ^19,31^, which are otherwise reported to localize to cytoplasmic puncta in response to other stresses ^18,20^, do not localize to puncta in response to HU stress. Similarly, Psp2 is a cytoplasmic RNA-binding protein with RGG motif implicated in translation ^17^, but it does not undergo a change in localization upon HU stress. These observations highlight the specific nature of change in localization of Scd6/Sbp1 upon HU stress. The reversibility of HU-induced Scd6 and Sbp1 puncta and their sensitivity to cycloheximide suggests that puncta-resident mRNAs may return to translation once the HU stress is removed.

The absence of Scd6 or Sbp1 improves the ability of cells to deal with HU stress. This is evident based on the results obtained from the growth curve and CFU experiments (Figures 2A-CC). We interpret these results to indicate that Scd6 is involved in repressing the mRNAs that encode proteins important for mounting a DNA damage response. At this stage, we do not know the identity of specific transcripts whose misregulation contributes to the improved ability of cells lacking Scd6 or Sbp1 to deal with HU stress.

SRS2 is a 3’-5’ DNA helicase whose expression levels are tightly controlled, which is evident from the observations that either overexpression or deletion of this gene causes sensitivity to genotoxins ^24,25^. Srs2 is a critical DNA damage response factorinvolved in dampening the DNA damage checkpoint, leading to genotoxin tolerance ^26^. Even though it has been established that Srs2 levels are tightly regulated in the cells ^24^, the regulator of this protein remains elusive. Our work identifies Scd6 as one such possible regulator of SRS2 by repressing its translation.

Our study provides mechanistic insights underlying the regulation of SRS2 by Scd6. We observe that Scd6 binds *SRS2* mRNA, and this interaction increases with HU stress. Interestingly, individual deletion of the LSm domain or RGG-motif compromises Scd6-*SRS2* interaction. We think that altered methylation levels of RGG-motif mediate the reduced interaction of the Scd6 LSm-domain deletion mutant with *SRS2*. The following observations support this: a) LSm-domain deletion mutant is hypermethylated as compared to full-length Scd6 (Figure 3D-F) and b) Methylated Scd6 interacts poorly with SRS2 as compared to unmethylated Scd6 (Figure 3H) . Direct cis-regulation of Scd6 AM by the Lsm domain is a unique observation that has not been reported for any methyltransferases. Results presented in Figure 3F-G indicate that the Lsm domain could either interact with Hmt1 or RGG-motif to negatively regulate Scd6 repression activity. The detailed basis of Lsm domain-mediated regulation of Scd6 methylation upon HU will be an interesting future direction.

Our results provide the first evidence for the role of human LSM14A as a translation regulator upon HU stress. Translation of multiple mRNAs is upregulated in LSM14A knockdown cells exposed to HU. We have validated changes in the translation status of two such mRNAs, *LIG4* and *RTEL1* (Figure 5D). RTEL1 is the functional homolog of SRS2, which, like SRS2, is an anti-recombinase and a negative regulator of HR-mediated DNA repair^32,33^. An interesting future direction would be to study the mechanism that allows translation repression and de-repression of these mRNAs by LSM14A. The repression mechanism of LSM14A is likely similar to Scd6, which targets eIF4G to repress translation ^15^. Although the observation that LSM14 is in physical proximity with eIF4G using proximity ligation assays supports this idea (Figure S6), further experiments in future will be required to confirm a conserved mechanism of repression by LSM14A.

LIG4 (DNA Ligase IV) is associated with XRCC4 to promote DNA double-strand break repair via the non-homologous end joining (NHEJ) pathway^34^. LSM14A knockdown increases the translation of LIG4 during HU stress (Figure 5D). This regulation has functional relevance as we observe that the NHEJ activity, as measured by plasmid integration assay, increases upon the knockdown of LSM14A under HU stress (Figure 5F). SRS2 has also been reported to be involved in the NHEJ pathway^35^. Therefore, like Scd6, LSM14A contributes to HU stress response by regulating the translation of a specific mRNA.

Like Scd6, Sbp1 also localizes to granules upon HU stress, and our studies indicate that this localization is also dependent on the RGG domain (Figure 1F). We observed that Scd6 and Sbp1 co-localize upon HU stress, and this localization is partly dependent on Scd6 (Figure S2C). But later results (Figure 2E) in this work indicate that both these proteins, even though have the same subcellular localization, have different, specific targets. An interesting future direction would be to understand the specific targets of Sbp1 that lead to HU stress tolerance and regulation of GSR. Overall, this study establishes RGG motif-containing proteins Scd6 and LSM14A as a regulator of genotoxic stress response by affecting translation of specific mRNA targets. Identifying the signalling mechanisms that enable genotoxic stress condition-specific repression activity of Scd6 and LSM14A would be a critical future direction.

## Methods and materials

### Yeast transformation

Strains were grown to 0.6 OD_600_ in complete media and pelleted down. The cells were washed once with water, followed by a single wash with 100 mM Lithium Acetate (LiAc). The cells were resuspended in 100 mM Lithium Acetate and aliquoted into 50 µL fractions. The cell suspension was then layered with 240 µL of 50 % PEG (v/v), 36 µL of 1 M LiAc, 25 µL of salmon sperm DNA (100 mg/ml) and and 100ng of the respective plasmid DNA and vortexed. The mixture was then incubated at 30 °C for 30 minutes, followed by 15 minutes at 42 °C. The cells were then pelleted, resuspended in 100 µL water and plated on synthetic defined media lacking Uracil (SD-Ura^-^) glucose agar plate. The plates were incubated at 30 °C for 2 days before colonies appeared.

### HU and Cycloheximide treatment

Yeast cultures were grown in 10 ml of SD-Ura^-^with glucose as the sugar, to 0.3-0.4 OD_600_ and split into two equal parts. The fractions were treated with 0.2 M HU or water (control) and incubated for 45 minutes (for Scd6) or 60 minutes (for Sbp1) at 30 °C with constant shaking. After incubation, 3 µL of 100 mg/ml Cycloheximide solution (dissolved in methanol) or methanol (control) was added to the HU-containing fractions. Cells were kept in the shaker for 5 minutes, followed by pelleting and live cell imaging.

### Yeast live cell imaging

In all cases for yeast live cell imaging, after the incubation period, the cells were immediately harvested (14000 rpm, 15 seconds), resuspended in 20 µL of media and spotted on a glass coverslip (No.1) and observed using live cell imaging. Yeast images were acquired using a Deltavision RT microscope system running softWoRx 3.5.1 software (Applied Precision, LLC), using an Olympus 100×, oil immersion 1.4 NA objective. The Green Fluorescent Protein (GFP) channel had 0.2 or 0.25 seconds of exposure (for Scd6GFP and Sbp1GFP, respectively) and 32 % transmittance. A minimum of 80 - 100 cells were observed for each experiment. Quantification was done as granules per cell. Statistical analysis was done using GraphPad Prism Version 7.0.

### Western Blotting

SDS Polyacrylamide gels were transferred onto Immobilon-P Transfer Membrane® (MERCK) using Transfer-Blot® Semi Dry apparatus (BIO-RAD). The transfer was done at 10 V for 1 hour. The membrane was then blocked with 5% skimmed milk, followed by washing and incubation with specific antibodies. For re-probing the blot with more than one antibody, the blot was stripped, blocked and reprobed.

| **Antibodies used in this study** | **Catalogue No.** |
| --- | --- |
| Anti-Sbp1 | Generated in lab |
| Anti-Pgk1 | Abcam, ab113687 |
| Anti-myc | Sigma, C3956 |
| Anti-monomethylation arginine | Cell Signalling Technologies, 8711S |
| Anti-His | Cell Signalling Technologies, 27E8 |
| Anti-Rabbit | Jackson ImmunoResearch Laboratories,<br>Code No. 111-035-003 |
| Anti-Mouse | Jackson ImmunoResearch Laboratories,<br>Code No. 115-035-003 |
| Monoclonal mouse LSM14A antibody | SantaCruz Biotechnology, sc-398552 |
| Rabbit EIF4G antibody | Cell Signalling Technologies, 2498 |
| Mouse EIF4E antibody | Cell Signalling Technologies, 9742 |
| Secondary mouse AlexaFluor 594 antibody | ThermoFisher , A32744 |

### Growth curve, Plating assay and Spot assay

For the growth curve, secondary yeast cultures were grown to the mid-log phase (0.4 to 0.6 OD_600_). The cells were then sub-cultured in fresh media to a final density of 0.1 OD_600_. The culture was then split into two parts, one treated with 0.2 M HU (final concentration) and the other with water (vehicle control). 200 μl of culture was aliquoted from each tube into 96 well-clear plates (Corning® Costar Clear Polystyrene 96-Well Plates, Cat ID: 3370). The plate was then incubated in a plate reader (Tecan Infinite 200 PRO) at 30 °C, with 40 seconds of orbital shaking (4 mm) every hour. Readings were recorded every hour for 24 hours. The data was analyzed and plotted using GraphPad Prism 5.

For spot assay, freshly grown yeast cells from the agar plate was resuspended in autoclaved deionized water and serially diluted to 10-fold dilutions after normalizing the first dilution to OD600 = 1. 10 μl of each dilution was spotted on an agar plate and incubated at 30 °C for 48-60 hours before imaging. For the plating assay, 100 μl of the 4th dilution, i.e., 10-4 OD600, was plated on a Control or 100 mM HU-containing agar plate. The plates were then incubated at 30 °C for 48-60 hours before counting the colony-forming units (CFUs).

### RNA Immunoprecipitation

300 ml of yeast cells expressing GFP-tagged Scd6, or Empty Vector (EV), were grown till OD_600_. The Scd6GFP culture was then split into two, where one was treated with 0.2 M HU and the other kept as an untreated control. The cultures were then pelleted and lysed in Lysis buffer [10 mM Tris, 150 mM NaCl, 1 X EDTA-free Protease inhibitor complex, RiboLock RNase Inhibitor, (Catalog No. EO0381)]) by bead beating. The lysate was spun at 5500 rpm for 5 mins. The supernatant was then transferred into a fresh MCT, and a small aliquot was removed for the isolation of total RNA. The rest of the lysate was diluted 1:1 with the dilution buffer, and 10 ul of GFP-TRAP Magnetic agarose beads (Cat. No. gtma) were added to a final volume of 1 ml. The pull-down samples were nutated at 4 °C for 90 minutes. The beads were then separated using a magnetic stand, followed by two washes with the lysis buffer. Finally, the beads were resuspended in 120 ul of lysis buffer. The pull-down efficiency was calculated by western blotting and probing the blot with an anti-GFP antibody. The rest of the pull-down was used for RNA isolation.

### RNA isolation and cDNA preparation

RNA isolation was done using Trizol (G Biosciences, Cat ID 786-652)-chloroform method. Briefly, the lysate/PD fractions were treated with 1 ml Trizol and 200 ul Chloroform. After vortexing, the mixture was spun at 14000 rpm for 20 minutes, followed by Isopropanol precipitation by flash freezing. It was spun at 150000 rpm at 4 °C for 30 minutes, followed by one 70% ethanol wash. The pellets were ly dried and resuspended in the required amount of DEPC-treated water. The RNA was then treated with DNAse (Thermo Scientific Cat ID: EN0521), followed by cDNA library preparation with 1ug of DNA-free RNA. OligodT was used for creating the libraries.

### Protein Purification and *In vitro* methylation

Recombinant N terminal His-tagged, and C terminal Flag-tagged Scd6 WT (His-Scd6-Flag) and domain deletion mutants were purified from *Escherichia coli* according to standard protocols using Ni-NTA agarose (G Biosciences Cat No.786-940) This was followed by a second purification with Flag beads (Sigma Aldrich Cat No.A2220) using standard purification protocols. N terminal His-tagged Hmt1 was purified using Ni-NTA agarose beads (G Biosciences Cat No. 786-940). Purified proteins were dialyzed with 20 mM Tris--Cl pH7.5, 100 mM NaCl, 10% glycerol and 1 mM DTT and purification was confirmed by SDS PAGE and Coomassie brilliant blue staining (Cat No. 32826). Purified His-Scd6-FLAG was methylated by taking 1:1 ratio of the protein and purified His-Hmt1 in methylation buffer containing 100 mM Tris–Cl pH8, 200 mM NaCl, 2 mM ethylenediaminetetraacetic acid (EDTA) and 1 mM Dithiothreitol (DTT) with or without 20 µM S-adenosyl methionine (SAM) (New England Biolabs) NEB; catalog no. B9003S). The reaction was incubated for 2 hours at 37 °C and used for downstream experiments. The methylation was confirmed by SDS-PAGE followed by western blotting using mono methyl arginine (MMA) antibody (Cell Signaling Technology) CST, catalog no. 8711; 1:1000 dilution)

### In vitro transcription (IVT) and Electromobility Shift Assay (EMSA)

The 5’ fragment of the SRS2 gene coding 184 bp of 5’UTR and 16 bps downstream the first codon of the ORF (200-mer template) was amplified from *BY4741* genomic DNA by PCR. 100 ng of PCR amplified, and column purified DNA template was transcribed using T7 polymerase (Thermo Cat ID. EP0111) in 100 ul reaction at 37 °C for 2 hours using manufacturer’s protocol followed by DNAse treatment (Thermo Scientific Cat ID: EN0521). The RNA was then purified using phenol-chloroform and stored in multiple aliquots for later use.

For EMSA, increasing concentrations of purified His-Scd6-Flag (0.7, 1.4, 2.1, 2.8, 3.4, 4.1 µM; unmethylated or methylated) were incubated with 0.94 µM of *in vitro* transcribed RNA in binding buffer [10 mM Tris (pH 8), 50 mM NaCl, 0.05% NP-40, 6% glycerol, 1 mM DTT and 2 μg/ml BSA] at 30 °C for 30 minutes. The reaction was then loaded on to 2% Agarose gel in 1 X MOPS buffer and the mobility shift visualized using ethidium bromide staining.

### Single-molecule Fluorescence *in situ* hybridization (smFISH)

smFISH protocol was adapted and modified from Tsanov *et. al*., 2016 ^36^ and Pizzinga *et. al*.,2019^37^. Briefly, WT and *Δsrs2* strains transformed with Scd6GFP plasmid were grown to mid-log phase (OD_600_ 0.5-0.8) and fixed with 3% paraformaldehyde for 45 min, at room temperature, in dark. To make smFISH probes, 200 pmol of an equimolar mix of SRS2-specific oligos (48 oligo pool, IDT) was annealed with 250 pmol of Cy5 labeled X-flap oligo in 1× NEBuffer 3. After fixation, cells were washed with buffer B (1.2 M sorbitol and 100 mM KHPO4, pH 7.5), followed by resuspension in spheroplasting buffer (Buffer B, 2 mM Ribonucleoside Vanadyl Complex, 0.2% β-mercaptoethanol, and 1 mg/ml lyticase) and incubated at 37°C shaker incubator for 15 min and stored in 70% ethanol at −20 °C . Subsequently, cells were hybridized with 40 pmol of smFISH probes in 100 µl of hybridization buffer (10 mg *E. coli* tRNA, 0.2 mM Ribonucleoside Vanadyl Complex, 200 µg/ml BSA, 10% dextran sulfate, 10% formamide, and 2× SSC). Cells were then washed in 10% formamide and 2× SSC. The washed cells were mixed with the mounting agent (Fluoromount-G™ Mounting Medium Cat ID. 00-4958-02) with DAPI, and spotted on the slide. The images were acquired using a Deltavision RT microscope system running softWoRx 3.5.1 software (Applied Precision, LLC), using an Olympus 100×, oil immersion 1.4 NA objective at required wavelengths. The images were processed and quantified using ImageJ software.

### Cell culture

The A375 and A2058 melanoma cell lines used in this study were purchased from the ATCC. Cancer cell lines were maintained at 37 °C and 5% CO_2_ in a humidified atmosphere and grown in DMEM:F12 or MEM growth media supplemented with 10% FBS, 2 mM glutamine, 50 u ml^−1^ penicillin and 50 mg ml^−1^ streptomycin (Gibco). RPE1 cells, used in Lsm14A-GFP live cell imaging, were grown in DMEM supplemented with 10% FBS and 2 mM L-glutamine. **LSM14A Live cell imaging**

For localization experiment with ectopically expressed LSM14A-GFP and LSM14AΔRGG1ΔRGG2-GFP. Cells were seeded on coverslips pre-coated with 1 μg/ml fibronectin (Sigma) and 20 μg/ml collagen (Sigma) 2 days before treatment with a genotoxic stress agent. After reaching confluency, cells were treated with either 10 mM hydroxyurea for 30 minutes or 10 µM of cisplatin for 6 hours. After the treatment, the coverslips were washed twice in PBS, and allowed to dry for 2-3 minutes. The cells were then fixed with 4% PFA in PBS for 10 minutes, followed by permeabilization for 10 min at room temperature in PBS containing 0.1% Tween-20 (PBST). Cells were washed with PBST, followed by 1X PBS. Cells were dried onto the cover slip before adding mounting media containing DAPI. This was followed by fixation onto a glass slide for confocal microscopy (Leica SPE)

### Polysome fractionation (Yeast)

WT cells expressing either EV or Scd6-GFP were grown till OD 0.8 and treated with 200 mM HU for 60 minutes, followed by 30mins of 0.1mg/ml cycloheximide treatment. The cells were lysed in lysis buffer (10mM Tris pH 7.4, 100mM NaCl, 30mM MgCl_2_, RNase inhibitor, cycloheximide and EDTA-free protease inhibitor complex), and pre-cleared lysate was loaded onto a 10–50% sucrose density gradient and centrifuged in an SW41 rotor (Beckman-Coulter) at 39,000 r.p.m for 2 h at 4 °C. The untranslated (till 80S fraction) and translated (polysome fractions) fractions were collected and pooled (Biocomp), and RNA was isolated from both fractions by TRIzol–chloroform method. Total RNA was also isolated from lysate. RNA samples were then treated with DNAse followed by DNA library preparation using random primers. RT-qPCR were performed using gene-specific primers.

### Polysome fractionation (mammalian cells)

To understand the translation control effect of LSM14A, A2058 cells were transfected with scrambled siRNA or siRNA specific to LSM14A cells. The cells were stressed with 10 mM hydroxyurea for 2 hours, followed by lysis, and polysome profiling. Sucrose density gradient centrifugation was used to separate the sub-polysomal and the polysomal ribosome fractions. Fifteen minutes before collection, cells were incubated at 37 °C with 100 mg/ml cycloheximide added to the culture medium. Next, cells were washed, scraped into ice-cold PBS supplemented with 100 mg/ml cycloheximide, centrifuged at 3,000 r.p.m. for 5 min and then collected into 400 ml of LSB buffer (20 mM Tris, pH 7.4, 100 mM NaCl, 3 mM MgCl_2_, 0.5 M sucrose, 2.4% Triton X-100, 1 mM DTT, 100 U ml RNasin and 100 mg/ml cycloheximide). After homogenization, 400 ml LSB buffer supplemented with 0.2% Triton X-100 and 0.25 M sucrose was added. Samples were centrifuged at 12,000g for 10 min at 4 °C. The resultant supernatant was adjusted to 5 M NaCl and 1 M MgCl_2_. The lysates were loaded onto a 15–50% sucrose density gradient and centrifuged in an SW41 rotor at 38,000 r.p.m. for 2 h at 4 °C. Polysome fractions were monitored and collected using a gradient fractionation system (Isco). Total RNA was extracted from the four heaviest fractions and the input samples using the TRIzol– chloroform method.

### Real-time PCR

RNA from each of the polysome fractions (16 fractions) was isolated using TRIzol LS reagent (TRIzol™ LS Reagent, Invitrogen, Cat ID 10296010) using the manufacturer’s protocol. The total isolated RNA was quantified, and 1 ug of RNA was used for complementary DNA synthesis. The cDNAs were diluted to 1:10 times from which equal volume (2 ul) cDNA were used along with SYBR green (TB Green Premix Ex Taq II (Tli RNase H Plus); TaKaRa, Cat ID RR820B) and primers (Bioserve, India) for validation of mRNA levels using quantitative real-time PCR (BioRad CFX96 Touch Real-Time PCR Detection System).

### RNA sequencing and bioinformatic analysis

RNA sequencing libraries were prepared from 500ng to 1µg of total RNA or mRNA-enriched from heavy polysome fractions using the Illumina TruSeq Stranded mRNA Library preparation kit which allows to perform a strand specific sequencing. Nanodrop spectrophotometer was used to assess purity of RNA based on absorbance ratios (260/280 and 260/230) and BioAnalyzer for RNA integrity (RIN>9). A first step of polyA+ selection using magnetic beads is done to focus sequencing on polyadenylated transcripts. After fragmentation, cDNA synthesis was performed and resulting fragments were used for dA-tailing followed by ligation of TruSeq indexed adapters. PCR amplification was finally achieved to generate the final barcoded cDNA libraries. The libraries were equimolarly pooled and subjected to qPCR quantification using the KAPA library quantification kit (Roche). Sequencing was carried out on the NovaSeq 6000 instrument from Illumina based on a 2*100 cycle mode (paired-end reads, 100 bases) to obtain around 30 million clusters (60 million raw paired-end reads) per sample. Finally, Fastq files were generated from raw sequencing data using bcl2fastq pipeline performing data demultiplexing based on index sequences.

RNA-seq data were analyzed with the Institut Curie RNA-seq Nextflow pipeline (Servant *et al*, 2023) (v3.1.4). Briefly, raw reads were first trimmed with Trimgalor. Reads were aligned on the human reference genome (hg19) using STAR (STAR_2.6.1a_08-27). Genes abundances were then estimated using STAR, and the Gencode v34 annotation. Differential analysis between conditions were done using the R package Xtail (Xiao *et al*, 2016) only on protein coding genes (Table S1).

### RNA interference

Cells were transfected with 20 nM of each siRNA against LSM14A (Dharmacon) and PRMT1 (Dharmacon) using Lipofectamine RNAiMAX Reagent (Life Technologies) following the supplier’s instructions.

### Proximity ligation assay

Interactions between EIF4E and EIF4G (EIF4E–EIF4G) or LSM14A and EIF4G (LSM14A-EIF4G) were detected by in situ proximity ligation assay (PLA) in A2058 melanoma cell lines untreated or treated with hydroxyurea. PLAs were performed on both fixed and permeabilized melanoma cells. The fixed, permeabilized cells were treated identically, and the PLA protocol was followed according to the manufacturer’s instructions (Olink Bioscience), with incubation of the primary antibodies at 4 °C overnight. After blocking, the antibodies were used at the following concentrations: for LSM14A (mouse, sc-398552, Santa Cruz Biotechnology, 1:200); for eIF4G (rabbit, 2498; Cell Signaling Technology, 1:200); for EIF4E (mouse, 9742, Cell Signaling Technology, 1:200). PLA minus and PLA plus probes (containing the secondary antibodies conjugated to oligonucleotides) were added and incubated for 1 h at 37 °C. More oligonucleotides were then added and allowed to hybridize to the PLA probes. Ligase was used to join the two hybridized oligonucleotides into a closed circle. The DNA was then amplified (with rolling circle amplification), and detection of the amplicons was carried out using the Brightfield detection kit for chromogenic development or using the Far-Red detection kit for fluorescence. Cell nuclei were stained with 4′,6-diamidino-2-phenylindole (DAPI). The sections were mounted with Olink Mounting Medium. The first results were visualized by confocal microscopy (Leica SPE), and the analysis was supported by Volocity software. To improve the sensitivity of the fluorescence detection, we next used a scanner (Olympus VS120) (magnification 20×; 2-ms exposure for the DAPI channel and 300-ms exposure for the Cy5 channel; 1 pixel = 0.32 μm), and the number of PLA signals per cell was counted (more than three fields) by semi-automated image analysis (ImageJ and OlyVIA).

## Data Availability

The datasets generated in this study have been deposited in the Gene Expression Omnibus repository (GEO) under accession numbers GSE274294 (token: qvutmiucrbydnwf)

## Acknowledgement

The authors thank Vagner and Rajyaguru labs for their critical inputs and suggestions during this work. The authors thank Won-Ki Huh (Seoul National University) for providing us with the *S. cerevisiae* Scd6-myc strain. We would also like to thank Martin Kupiec for kindly gifting pRS316-SRS2 plasmid and Professor Alison Bertuch for Yku70 and Yku80 constructs. We thank Poornima Gopalakrishna for creating the RGG-deletion mutants of LSM14A.

## Supplementary Figures

**Figure S1.**
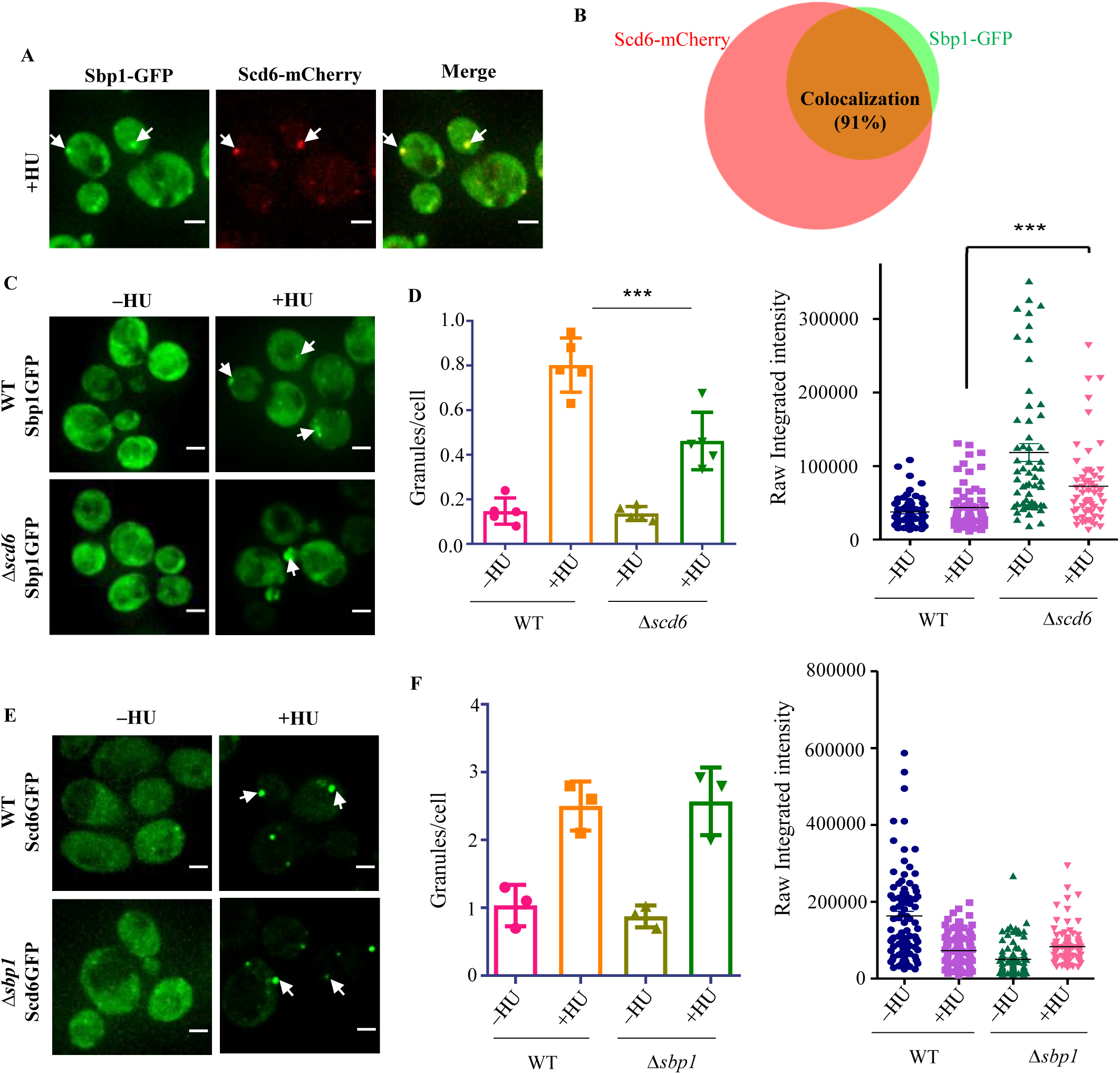
Sbp1 localizes to granules in an Scd6-dependant manner. (A) Live cell imaging showing colocalization of Sbp1 and Scd6 granules in HU- treated cells. (B) Venn diagram showing colocalization of Scd6-mCherry and Sbp1-GFP granules (Red denotes Scd6-mCherry granules and green indicates Sbp1-GFP granules. Brown colour indicates colocalizing granules). ∼90% of Sbp1 granules colocalize with Scd6 granules upon 60 mins of HU treatment (n=3; ∼300 cells were counted). Percentage colocalization was calculated by dividing total co-localizing foci by the total number of Sbp1 foci (C) and (E). Live cell imaging showing localization of Sbp1GFP and Scd6-GFP in wild type (WT) and *Δscd6* or *Δsbp1* strains, respectively. (D) and (F) Quantification of (C) and (E) as granules/cell and raw integrated GFP intensity (n=3;∼400 cells were counted; 100 cells were used for intensity calculations. Error bars indicate standard error of mean. Statistical significance was calculated using unpaired t-test. Scale bar denotes 2 μm.

**Figure S2.**
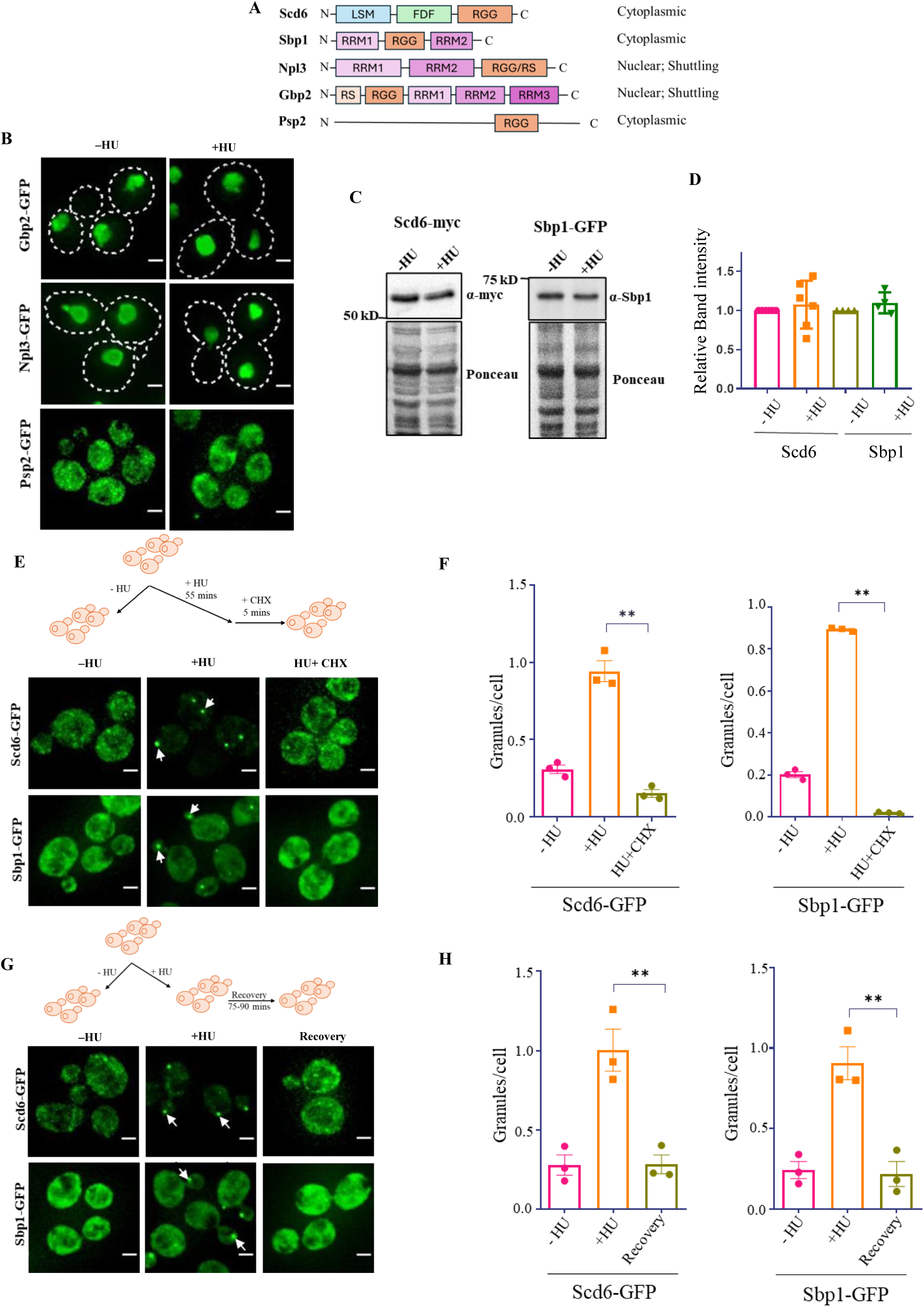
Scd6 and Sbp1 form reversible, RNA containing granules upon HU treatment. (A) Schematic representation of the domain organization of Scd6, Sbp1, Npl3, Gbp2 and Psp2 (B) Gbp2, Npl3, and Psp2, another set of RGG motif-containing RNA binding proteins, do not show a change in localization to granules even after 120 mins of HU treatment. Dotted lines represent cell boundaries. (C) Western blot showing protein levels of tagged Scd6 and Sbp1 proteins upon HU treatment (45 minutes and 60 minutes respectively). Ponceau stained blot was used as loading control (D) Quantification of Scd6-myc and Sbp1-GFP protein levels shown in C. (E) Live cell image showing the change in localization of endogenously tagged Scd6-GFP and Sbp1-GFP after 5 minutes of 0.1 mg/ml CHX treatment. (F) Quantification of granules as granules per cell (Left panel shows granule count for Scd6 and right panel for Sbp1). (G) Live cell imaging showing Scd6 and Sbp1 localization after incubating in non–HU-containing media (Recovery period is 75 min for Scd6 and 90 minutes for Sbp1). (H) Quantification of granules in terms of granules per cell (Left panel shows change in granules for Scd6 and the right panel shows the same for Sbp1. n=3; ∼300 cells were counted for analysis for all experiments. Statistical significance was calculated using unpaired t-test. Error bars indicate standard error of mean (SEM). Scale bar denotes 2 μm.

**Figure S3.**
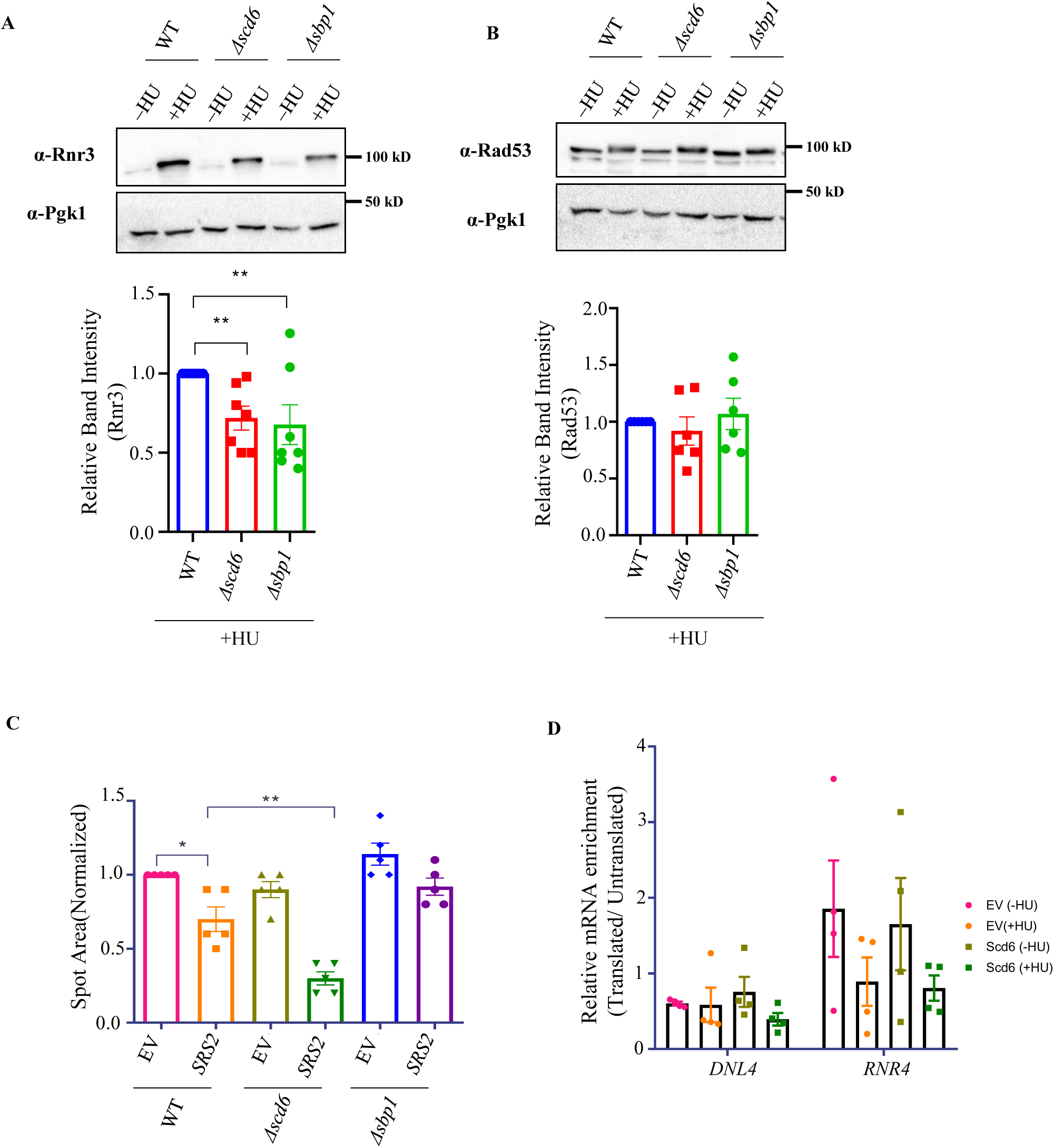
Scd6 modulates cellular tolerance to HU. (A) and (B) Western blots probed for Rnr3 and Rad53 protein levels using protein specific antibody, and their respective quantification (below) for cells treated with 200 mM HU for 90 minutes. (C) Quantification of growth assay (in Figure 2E) by measuring the intensity of the second spot and normalizing the values of spots on HU plate to the respective control values for each strain (n=5). (D) Quantification of *DNL4* and *RNR4* mRNA in the polysome fractions plotted as relative log_2_-Fold change ratio of Translated/Untranslated fractions (n=4) using SRS2 specific primers. Error bars indicate standard error of mean and statistical significance was calculated using unpaired t-test.

**Figure S4.**
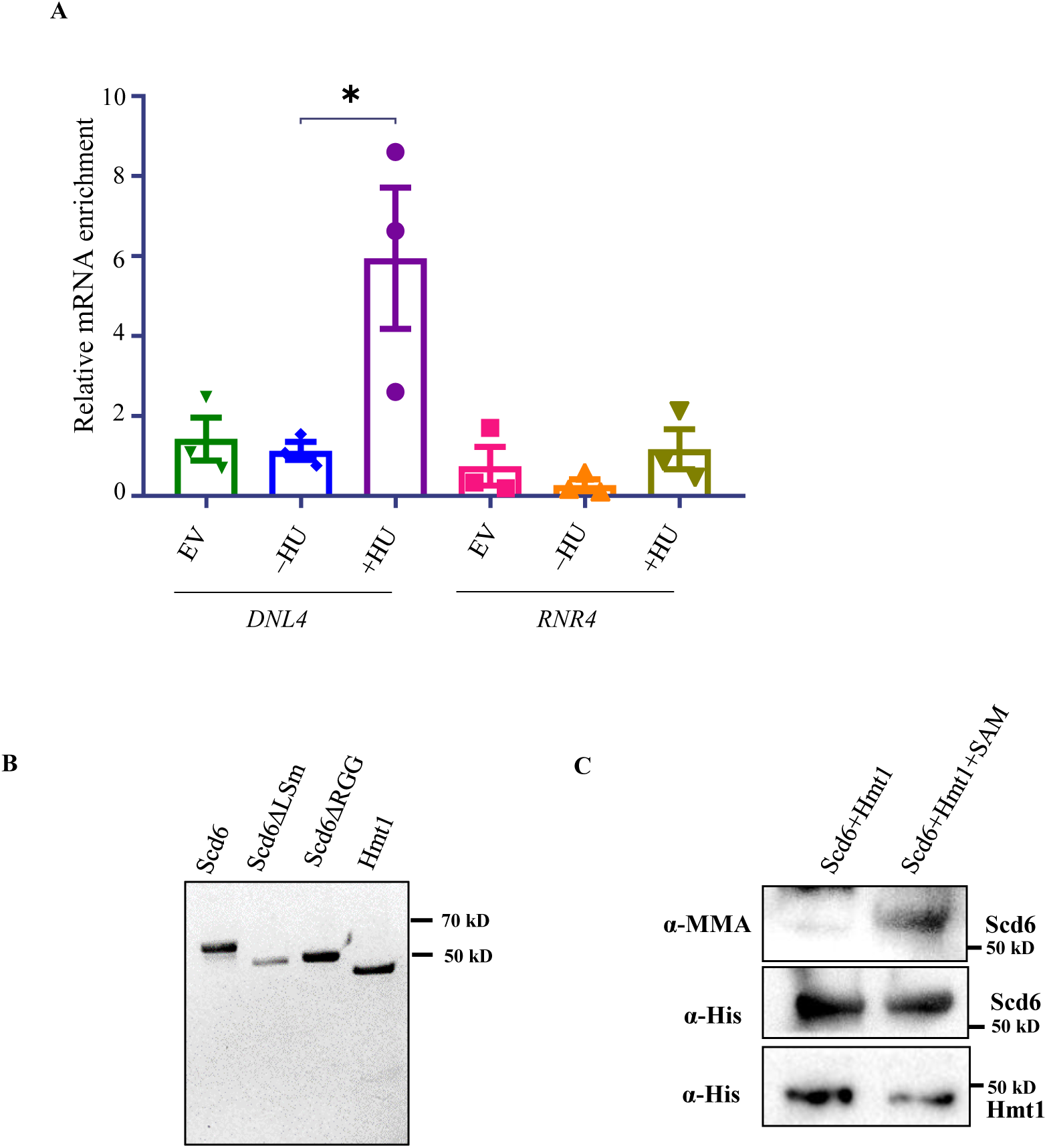
Scd6 binds to specific mRNA. (A).Quantification of mRNA enrichment in RNA immunoprecipitation with *DNL4* and *RNR4* specific primers. Statistical significance was calculated using unpaired t-test. Error bars indicate standard error of mean. (B) Coomassie Brilliant Blue stained gel of purified His-Scd6-FLAG (HSF), HSF-ΔLSm, HSF-ΔRGG and His-Hmt1 (C) Western blot showing methylation of purified HSF in the presence of His-Hmt1 and 1 mM S-adenosyl methionine (SAM).

**Figure S5.**
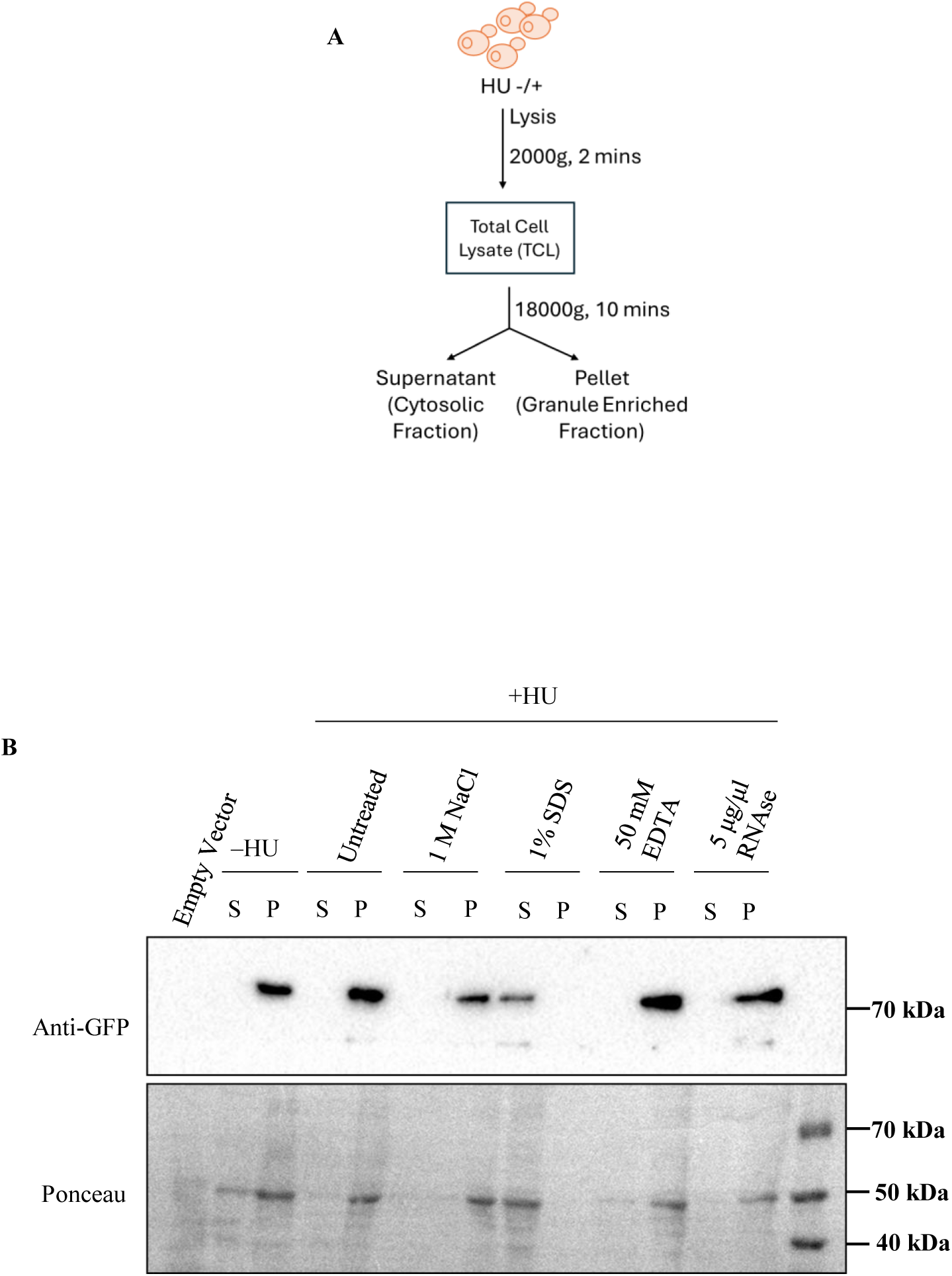
Scd6 localizes to stress granule cores upon HU stress. (A) Schematic showing workflow for granule enrichment from cells expressing Scd6-GFP in the presence or absence of 200 mM HU. (B) Western blot showing partitioning of granule enriched Scd6GFP into soluble (S) or pellet (P) fraction upon treatment with 1M NaCl, 1% SDS, 50 mM EDTA or 5 μg/μl RNAse.

**Figure S6.**
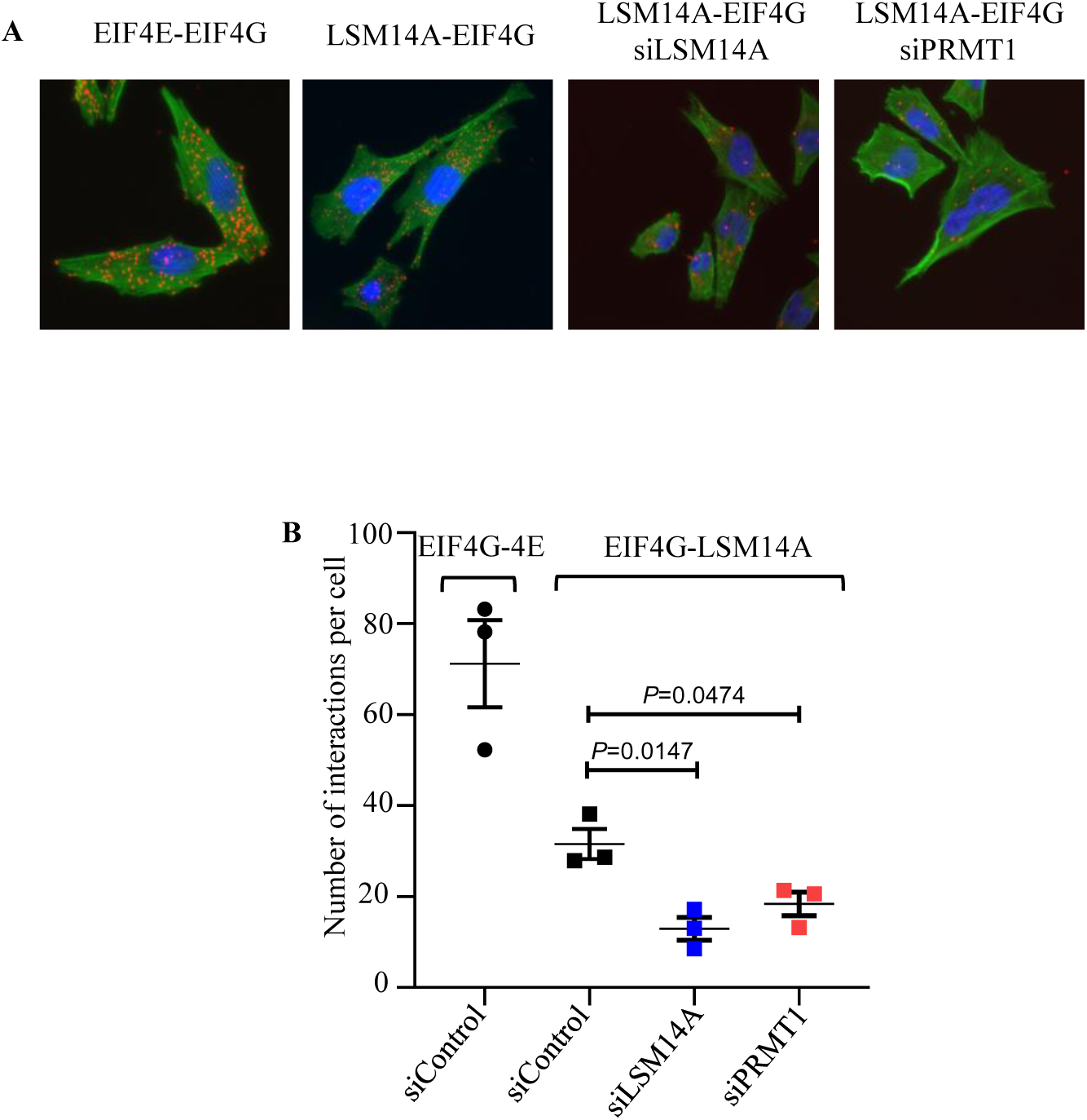
Proximity ligation assay reveals interaction of LSM14A and EIF4G. (A) Proximity ligation assay to show the interaction between EIF4G-EIF4E (positive control) and LSM14A-EIF4G. The interaction between EIF4G and LSM14A is reduced when the expression of LSM14A was downregulated using small-interfering RNA against LSM14A. Similarly, the interaction of LSM14A and EIF4G reduced significantly when PRMT1 was downregulated in A2058 cells. (B) Quantitation of proximity ligation assay data presented in A

## References

1. Marnett, L. Endogenous DNA damage and mutation. Trends Genet. 17, 214–221 (2001).

2. Mohanan, G., Das, A. & Rajyaguru, P. I. Genotoxic stress response: What is the role of cytoplasmic mRNA fate? BioEssays 43, 2000311 (2021).

3. Thandapani, P., O’Connor, T. R., Bailey, T. L. & Richard, S. Defining the RGG/RG Motif. Mol. Cell 50, 613–623 (2013).

4. Chowdhury, M. N. & Jin, H. The RGG motif proteins: Interactions, functions, and regulations. WIREs RNA (2022) doi:10.1002/wrna.1748.

5. Chong, P. A., Vernon, R. M. & Forman-Kay, J. D. RGG/RG Motif Regions in RNA Binding and Phase Separation. J. Mol. Biol. 430, 4650–4665 (2018).

6. Buchan, J. R. & Parker, R. Eukaryotic Stress Granules: The Ins and Outs of Translation. Mol. Cell 36, 932–941 (2009).

7. Rajyaguru, P., She, M. & Parker, R. Scd6 Targets eIF4G to Repress Translation: RGG Motif Proteins as a Class of eIF4G-Binding Proteins. Mol. Cell 45, 244–254 (2012).

8. Bhatter, N. et al. Arginine methylation augments Sbp1 function in translation repression and decapping. FEBS J. 286, 4693–4708 (2019).

9. Poornima, G., Shah, S., Vignesh, V., Parker, R. & Rajyaguru, P. I. Arginine methylation promotes translation repression activity of eIF4G-binding protein, Scd6. Nucleic Acids Res. 44, 9358–9368 (2016).

10. Roy, R., Das, G., Kuttanda, I. A., Bhatter, N. & Rajyaguru, P. I. Low complexity RGG-motif sequence is required for Processing body (P-body) disassembly. Nat. Commun. 2022 131 **13**, 1–13 (2022).

11. Yang, W. H., Jiang, H. Y., Gulick, T., Bloch, K. D. & Bloch, D. B. RNA-associated protein 55 (RAP55) localizes to mRNA processing bodies and stress granules. RNA (2006) doi:10.1261/rna.2302706.

12. Roy, D. & Rajyaguru, P. I. Suppressor of clathrin deficiency (Scd6)—An emerging RGG-motif translation repressor. Wiley Interdiscip. Rev. RNA 9, e1479 (2018).

13. Brandmann, T. et al. Molecular architecture of LSM14 interactions involved in the assembly of mRNA silencing complexes. EMBO J. 37, (2018).

14. Tanaka, K. J. et al. RAP55, a Cytoplasmic mRNP Component, Represses Translation in Xenopus Oocytes. J. Biol. Chem. 281, 40096–40106 (2006).

15. Li, Y. et al. LSm14A is a processing body-associated sensor of viral nucleic acids that initiates cellular antiviral response in the early phase of viral infection. Proc. Natl. Acad. Sci. 109, 11770–11775 (2012).

16. Tkach, J. M. et al. Dissecting DNA damage response pathways by analysing protein localization and abundance changes during DNA replication stress. Nat. Cell Biol. 14, 966–976 (2012).

17. Yin, Z. et al. Psp2, a novel regulator of autophagy that promotes autophagy-related protein translation. Cell Res. (2019) doi:10.1038/s41422-019-0246-4.

18. Poornima, G. et al. RGG-motif containing mRNA export factor Gbp2 acts as a translation repressor. RNA Biol. (2021) doi:10.1080/15476286.2021.1910403.

19. Flach, J. et al. A yeast RNA-binding protein shuttles between the nucleus and the cytoplasm. Mol. Cell. Biol. (1994) doi:10.1128/mcb.14.12.8399.

20. Windgassen, M. et al. Yeast Shuttling SR Proteins Npl3p, Gbp2p, and Hrb1p Are Part of the Translating mRNPs, and Npl3p Can Function as a Translational Repressor. Mol. Cell. Biol. 24, 10479–10491 (2004).

21. Brengues, M., Teixeira, D. & Parker, R. Movement of eukaryotic mRNAs between polysomes and cytoplasmic processing bodies. Science 310, 486–489 (2005).

22. Singh, A. & Xu, Y.-J. The cell killing mechanisms of hydroxyurea. Genes (Basel). 7, 99 (2016).

23. Marini, V. & Krejci, L. Srs2: The “Odd-Job Man” in DNA repair. DNA Repair (Amst). 9, 268–275 (2010).

24. Bronstein, A., Bramson, S., Shemesh, K., Liefshitz, B. & Kupiec, M. Tight Regulation of Srs2 Helicase Activity Is Crucial for Proper Functioning of DNA Repair Mechanisms. G3 Genes|Genomes|Genetics 8, 1615–1626 (2018).

25. Alex, B., Lihi, G., Gilad, G., Elisa, A.-P. & Martin, K. The Main Role of Srs2 in DNA Repair Depends on Its Helicase Activity, Rather than on Its Interactions with PCNA or Rad51. MBio 9, 10.1128/mbio.01192-18 (2018).

26. Dhingra, N. et al. The Srs2 helicase dampens DNA damage checkpoint by recycling RPA from chromatin. Proc. Natl. Acad. Sci. 118, (2021).

27. Wheeler, J. R., Jain, S., Khong, A. & Parker, R. Isolation of yeast and mammalian stress granule cores. Methods 126, 12–17 (2017).

28. Singh, A. & Xu, Y. J. The cell killing mechanisms of hydroxyurea. Genes (Basel). 7, (2016).

29. Koç, A., Wheeler, L. J., Mathews, C. K. & Merrill, G. F. Hydroxyurea Arrests DNA Replication by a Mechanism that Preserves Basal dNTP Pools. J. Biol. Chem. 279, 223–230 (2004).

30. Saintigny, Y. Characterization of homologous recombination induced by replication inhibition in mammalian cells. EMBO J. 20, 3861–3870 (2001).

31. Windgassen, M. & Krebber, H. Identification of Gbp2 as a novel poly(A) + RNA-binding protein involved in the cytoplasmic delivery of messenger RNAs in yeast. EMBO Rep. 4, 278–283 (2003).

32. Frizzell, A. et al. RTEL1 Inhibits Trinucleotide Repeat Expansions and Fragility. Cell Rep. 6, 827–835 (2014).

33. Barber, L. J. et al. RTEL1 Maintains Genomic Stability by Suppressing Homologous Recombination. Cell 135, 261–271 (2008).

34. Grawunder, U., Zimmer, D., Kulesza, P. & Lieber, M. R. Requirement for an Interaction of XRCC4 with DNA Ligase IV for Wild-type V(D)J Recombination and DNA Double-strand Break Repairin Vivo. J. Biol. Chem. 273, 24708–24714 (1998).

35. Hegde, V. Requirement for the SRS2 DNA helicase gene in non-homologous end joining in yeast. Nucleic Acids Res. 28, 2779–2783 (2000).

36. Tsanov, N. et al. smiFISH and FISH-quant – a flexible single RNA detection approach with super-resolution capability. Nucleic Acids Res. 44, e165–e165 (2016).

37. Pizzinga, M. et al. Translation factor mRNA granules direct protein synthetic capacity to regions of polarized growth. J. Cell Biol. 218, 1564–1581 (2019).

